# False-Positive IRESes from *Hoxa9* and other genes resulting from errors in mammalian 5’ UTR annotations

**DOI:** 10.1101/2022.02.10.479744

**Authors:** Christina Akirtava, Gemma E. May, C. Joel McManus

**Author notes:** Correspondence to C. Joel McManus. These authors contributed equally to this study (Christina Akirtava, Gemma E. May). **Author Contributions:** C.A., G.E.M., and C. J. M. designed research. C.A. and G.E.M. performed research and analyzed data. C.J.M. wrote the manuscript, with input from all authors.

## Abstract

Hyperconserved genomic sequences have great promise for understanding core biological processes. It has been recently proposed that scores of hyperconserved transcript leaders (hTLs) encode Internal Ribosome Entry Sites (IRESes) that drive cap-independent translation in part via interactions with ribosome expansion segments. However, the direct functional significance of such interactions has not yet been definitively demonstrated. We provide evidence that the putative IRESes previously reported in Hox gene hTLs are rarely included in transcript leaders. Instead, these regions function independently as transcriptional promoters. In addition, we find the proposed RNA structure of the putative *Hoxa9* IRES is not conserved. Instead, sequences previously shown to be essential for putative IRES activity encode a hyperconserved transcription factor binding site (E-box) that contributes to its promoter activity by binding to the transcription factors *USF1* and *USF2*. Similar E-box sequences enhance the promoter activities of other putative *Hoxa* gene IRESes. Moreover, we provide evidence that the vast majority of hTLs with putative IRES activity overlap transcriptional promoters, enhancers, and 3’ splice sites that are most likely responsible for their reported IRES activities. These results argue strongly against recently reported widespread IRES-like activities from hTLs and contradict proposed interactions between ribosomal expansion segment ES9S and putative IRESes. Furthermore, our work underscores the importance of accurate transcript annotations, controls in bicistronic reporter assays, and the power of synthesizing publicly available data from multiple sources.

## Introduction

As a critical step in gene expression, the translation of mRNA into protein is highly regulated. Eukaryotic translation is primarily controlled at the initiation stage, in which ribosomes identify start codons and begin synthesizing protein (1, 2). During proliferative growth, most mRNA translation initiates through a cap-dependent mechanism in which the 5’ 7-mG interacts with initiation factors to recruit a pre-initiation complex (PIC) comprised of the 40S small ribosomal subunit and multiple initiation factors. Once recruited, PICs scan directionally 5’ to 3’ until a start codon is recognized, the large ribosomal subunit is recruited, and translation commences. Under stress conditions, this cap-dependent translation is largely repressed due to inactivation of initiation factors. In such circumstances, ribosomes can be recruited to mRNA through cap-independent mechanisms, including Internal Ribosome Entry Sites (IRESes). IRESes are often found in viruses, as these pathogens often suppress cap-dependent translation of cellular RNAs to commandeer ribosomes for viral protein synthesis. IRESes have also been reported in cellular mRNA, though their roles in translation remain controversial (1, 2).

Several studies have coalesced on a surprising model in which hyperconserved transcript leaders include IRES-like sequences that drive cap-independent translation in specific cell types during development. These IRES-like elements were first proposed for mammalian *Hoxa* genes, based on the observation that the annotated mouse transcript leaders from several *Hoxa* genes drove expression in bicistronic luciferase assays, a classic test for cap-independent translation (3). It has also been proposed that ribosome expansion segment 9S (ES9S), a stem loop that protrudes from the ribosome, binds to a structured stem loop in the *Hoxa9* IRES-like sequence to recruit ribosomes to the *Hoxa9* transcript (4). Mammalian ES9S was also shown to bind G-rich motifs found in many mRNAs *in vitro*, which was proposed to drive cap-independent translation of many cellular transcripts (5).

The possibility of widespread cap-independent translation driven by interactions between IRESes and expansion segments is tantalizing. However, many previously reported IRESes in cellular mRNA have been fraught with controversy (6–8), especially when the sole evidence for such IRESes comes from bicistronic reporter assays. In these assays, a test IRES sequence is cloned between two luciferase open reading frames, with the expectation that the downstream luciferase will only be expressed if the test sequence is an IRES. However, this assay is widely known to produce false positives resulting from monocistronic transcripts from transcriptional promoters or cryptic splicing in the IRES test sequence (9–14)(Figure S1). The bicistronic plasmid used in these studies (pRF) also has cryptic upstream promoters that generate unexpected monocistronic transcripts, which further complicates the interpretation of assay results (15)(Figure S1), It has also been noted that the Hox genes likely have much shorter transcript leaders than those used in bicistronic IRES assays (7). Furthermore, previous RNAi control experiments suggested that putative *Hoxa* gene IRESes have independent promoter activity. While siRNA targeting the upstream *Rluc* eliminated *Rluc* expression,∼ 30% of *Fluc* expression resisted RNAi, indicating substantial monocistronic Fluc transcripts (3). However, the authors of the study inexplicably drew the opposite conclusion.

The proposed functional interaction between the *Hoxa9* P4 stemloop and ribosome ES9S is also problematic. Previous work found sequences complementary to human ES9S did not support IRES activity (16). In addition, the proposed IRES RNA structures are inconsistent with functional assays. A cryo-EM structure model of this interaction appears to show the helices oriented as kissing stem loops. This structure most likely involves base pairing between nucleotides in the G-rich P4 loop with the C-rich loop of ES9S (Figure S2). However, mutations to the G-rich loop of P4 did not disrupt its apparent IRES activity (4, 5), suggesting that this proposed interaction is dispensable. Although some mutations that disrupt the P4 stem greatly reduced apparent IRES activity, compensatory mutations to restore P4 base pairing did not restore IRES function (4). Finally, a deletion of the 5’ half of the P4 stem loop did not disrupt IRES-like activity in the bicistronic reporter. As such, the authors could not rationalize how the P4 / hES9S interaction visualized by cryo-EM related to bicistronic reporter expression. Together, these observations cast doubt on the model that ES9S binds to the *Hoxa9* P4 stem loop to drive cap-independent translation.

Recently, a high-throughput analysis defined a set of 589 hyperconserved transcript leaders (hTLs) with strong enrichment for genes involved in mammalian development (17). Hundreds of these hTLs were tested for IRES-like activities in bicistronic reporter assays and thirty-seven percent (90 / 241) drove substantial expression of the downstream luciferase cistron, suggesting that hTLs may frequently encode IRES-like functional elements. However, the possibility that these putative IRES activities may instead reflect functional promoter elements or cryptic 3’ splices sites was not directly investigated. While the authors showed *Fluc* / *Rluc* protein and RNA ratios were not strongly correlated, this could result from variance in luciferase and RT-qPCR measurements (18, 19), especially considering that the pRF reporter plasmid expresses cryptic transcripts that would also be amplified (15). Indeed, previous work has cautioned against using RT-qPCR to normalize bicistronic reporter assays (20). Consequently, though they provide an alluring model for new modes of translational control during mammalian development, the authenticity of hTL IRES-like elements has not been conclusively established.

In this work, we investigated the possibility that putative IRES-like elements in mammalian hTLs instead encode transcriptional promoters and 3’ splice sites. We show that the putative *Hoxa9* IRES shows no signs of structural conservation and does not appear to be expressed at biologically relevant levels. Instead, sequences encoding putative IRESes from mouse *Hoxa* genes act as independent promoters. In addition, we demonstrate that a sequence previously identified as essential for *Hoxa9* IRES activity is a classical “E-box” site recognized by bHLH transcription factors. Putative IRESes from other *Hoxa* genes similarly have conserved E-box motifs that contribute to their promoter activities. Furthermore, the proposed IRES-like elements in the transcript leaders of *Chrdl1*, *Cnot3*, *Cryab* and *Slc25a14* also have strong promoter activities. We also find putative hTLs frequently overlap other functional elements, including protein coding sequences, which could explain their conservation. Finally, we show that recently proposed IRES-like hTLs are overwhelmingly further enriched in annotated promoters, 3’ splice sites, and internal transcription initiation, and these elements can be used to accurately predict their reported IRES-like activities.

## Results

We first investigated the putative IRES region of *Hoxa9*, which has been called the paradigmatic example hyperconserved transcript leader (17). Many IRESes, including viral IRESes, fold into complex functional RNA structures. Previous studies reported a complex secondary structure for the *Hoxa9* IRES based on SHAPE probing (3). Two RNA base pairing regions (P3a and P4) were required for IRES activity (3, 4). The P4 stem-loop was later shown to interact with ribosomal expansion segment ES9S *in vitro* (4), yet the P4 structure was not required for bicistronic reporter activity (4). Thus, the functional significance of the P4 region remains unclear. To further investigate the importance of the *Hoxa9* IRES structure, we examined its conservation. The putative IRES regions of both mouse and zebrafish *Hoxa9* were previously shown to drive bicistronic reporter expression in mouse cell culture (3). However, the predicted RNA structures of mouse, human and zebrafish *Hoxa9* IRES regions differ substantially, such that the P3a domain is not predicted to form in the homologous human sequence, and neither P3a or P4 are likely to form in zebrafish (Figures 1A & B). Furthermore, we found no evidence of significant structural covariation in RNA sequence alignments from 230 mammals (21) using Infernal (22) and R-scape (23) (Figure 1C), despite having enough statistical power to detect such pairing (24) (Table S1). These results indicate that the proposed RNA structure of the *Hoxa9* IRES region, including domains previously reported to be essential for IRES activity, is not evolutionarily constrained.

**Figure 1.**
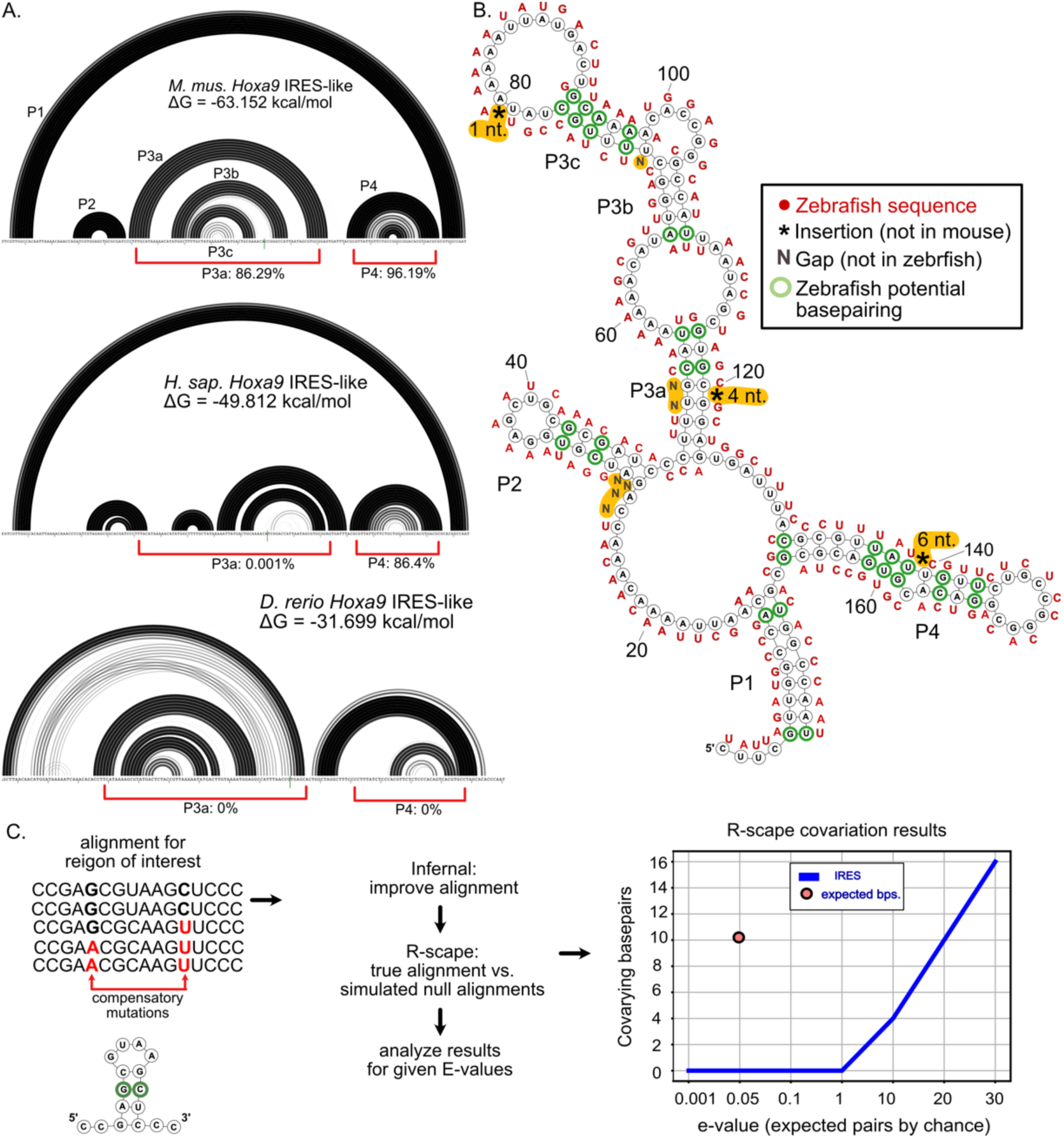
The proposed structure of the putative *Hoxa9* IRES element is not conserved. (A) Rainbow graphs depict the probability of base pairing for the *Hoxa9* IRES-like region from mouse (top), human (middle) and zebrafish (bottom). Pairing regions (stems) are numbered as in (3). Base pairing probabilities were determined using RNAstructure. The mouse model is highly consistent with the published model (3). Red brackets indicate the frequency of P3a and P4 helix formation in 10,000 predicted suboptimal structures. Human and mouse *Hoxa9* share P1 and P4, but lack P3, which was reported to be essential for IRES activity (3). Zebrafish *Hoxa9* does not share any structural similarity with mammalian homologs, despite driving bicistronic reporter activity (3). (B) Secondary structure model of mouse *Hoxa9* putative IRES region (3). Corresponding zebrafish sequences are shown in red. Most proposed base pairs are not conserved. Zebrafish has insertions (asterisks) and deletions (Ns) in the critical P3 and P4 elements. (C) Results of R-scape analysis of mutual information for the mouse *Hoxa9* putative IRES region using alignments from 208 mammals and 23 other vertebrates. The number of covarying sites (y axis) is given for different e-value cutoffs (x axis). Although the alignment has the power to detect ∼10 compensatory pairs (red point; Table S1), covarying base pairs are less common than expected by chance in the IRES-like element (blue line).

To drive cap-independent translation, the mouse *Hoxa9* IRES must be included in its TL. Previous work noted that this may not be the case (7). Our evaluation of public RNA-seq data indicates that the annotated TL of mouse *Hoxa9* used in previous studies shows little evidence of transcription in mouse tissues (Figure 2, Figure S3). In nearly all tissues analyzed, including embryonic neural tube, the tissue from which the IRES was first reported (3), ENCODE RNA-seq data show negligible levels of transcribed RNA in the upstream region of the annotated 5’ UTR. Instead, transcript levels sharply increase close to the *Hoxa9* start codon, immediately downstream of a strong transcription start site (TSS) annotated in the refTSS database. These short 5’ UTR isoforms are also supported by ENCODE long-read RNA-seq data from developing embryos (Figure 2). Although the extended TL is partially supported by one long read, this appears to be an unspliced intron from *Hoxa10*/*Hoxa9* or *Mir196b*/*Hoxa9* fusion transcripts (Figure 2). A very similar set of isoforms is supported by human RNA-seq data (Figure S3) (25–1. 27) and also suggest the putative IRES region is excluded from translating mRNA (Figure S4) (28). Thus, the putative *Hoxa9* IRES is almost completely excluded from mouse and human *Hoxa9* transcripts.

**Figure 2.**
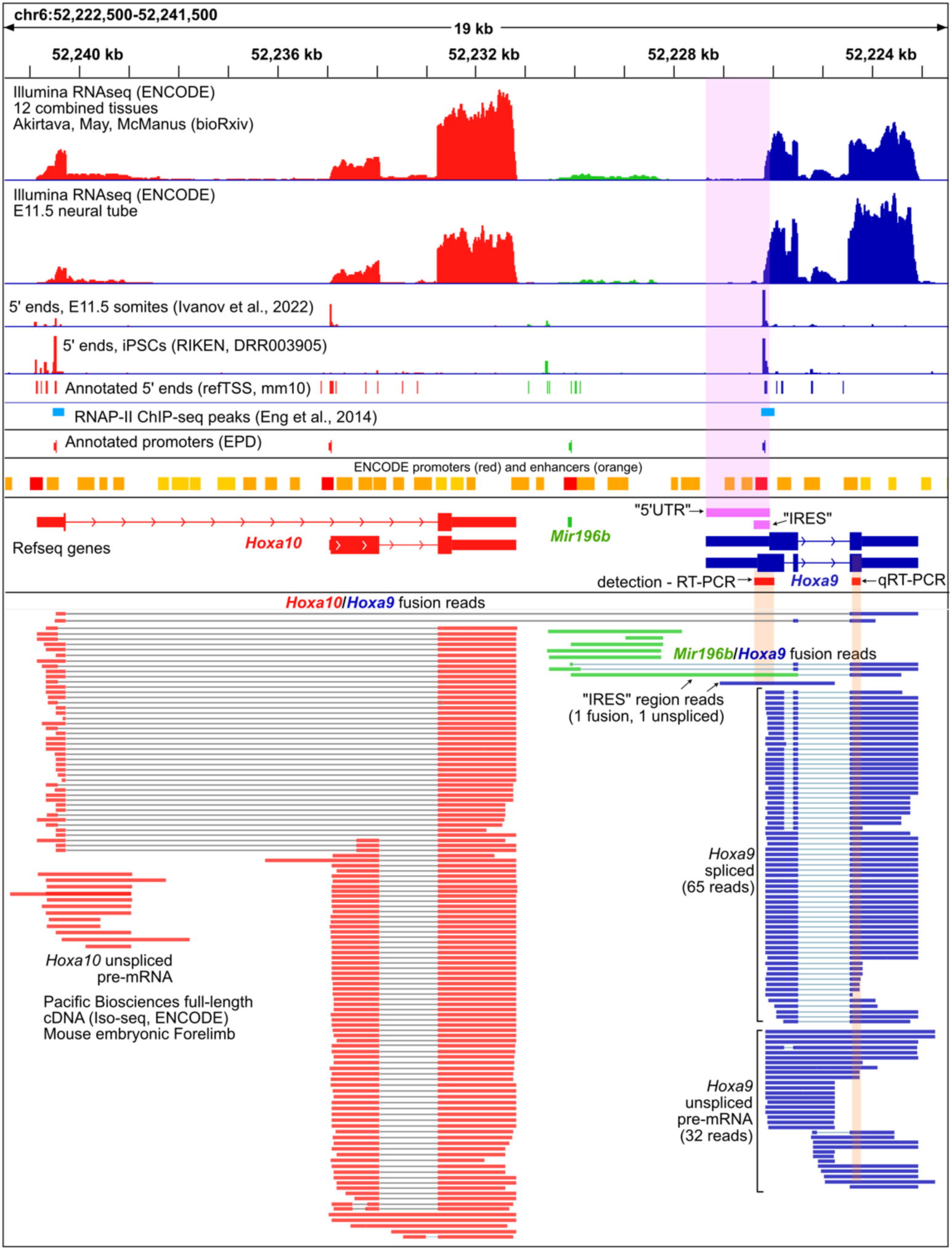
The putative extended 5’ UTR and IRES regions of mouse *Hoxa9* are not expressed at biologically meaningful levels. A genome browser view showing Refseq annotations of the mouse *Hoxa10* / *Mir196b* / *Hoxa9* in red, green, and blue, respectively, is shown. The putative TL(5’UTR) and IRES regions are shown in pink. Promoters from the Eukaryotic Promoter Database (EPD) and the ENCODE project are shown above the Refseq gene models. Illumina short-read (upper) and PacBio full-length (lower) RNA-seq data from the ENCODE consortium show negligible levels of RNA over the putative 5’ UTR and IRES regions. Similarly, two 5’ CAGE-seq studies (Ivanov et al., 2022; Abugessaia et al., 2019) show no transcription start sites (TSS) at the putative 5’ UTR, and a strong TSS peak downstream of the putative IRES. Chip-seq peaks show RNAPII is found immediately downstream of EPD *Hoxa9* and *Hoxa10* promoters in mouse embryonic forelimbs (Eng et al., 2014). PacBio RNA-seq detects *Hoxa9*/*a10* and *Hoxa9*/*Mir196b* fusion transcripts. Regions corresponding to the PCR amplicons used previously to detect (RT-PCR) the putative *Hoxa9* IRES and quantify (RT-qPCR) *Hoxa9* mRNA are shown in red. The *Hoxa9* RT-qPCR amplicon used for expression and polysome analysis is not specific to spliced *Hoxa9* mRNA, and can amplify fusion transcripts, unspliced transcripts, and truncated transcripts initiating at RefTSS annotated start sites within *Hoxa9* introns.

As the putative *Hoxa9* IRES appears to be part of an intron from rare fusion transcripts, we examined an alternative hypothesis that the reported UTR region encodes functional DNA elements. Manual examination revealed the extended TL region overlapped two enhancers and a promoter annotated by the ENCODE consortium and the Eukaryotic Promoter Database (EPD) (21, 22)(Figure 2). In addition, public ChIP-seq data from mouse embryos show RNAPII peaks just after the EPD promoter (29), consistent with promoter proximal pausing (30, 31) (Figure 2). We tested this region for promoter activity in mouse C3H/10T1/2 embryonic mesenchymal cells, which were previously used to study the *Hoxa9* IRES (3). To most directly compare our results to previous IRES studies, we used a modified bicistronic reporter plasmid lacking the upstream SV40 promoter (pRF-ΔSV40) and cloned putative *Hoxa9* promoter regions between *Rluc* and *Fluc* (Figure 3A). Strikingly, both the putative extended TL and putative IRES regions of mouse and human *Hoxa9* drove expression of Firefly luciferase. Additionally, this expression was absent in reporters in which the putative IRES region was reversed (Figure 3B). Finally, we used 5’ RACE to map *F. luc* TSSs used in the *Hoxa9* reporter and found they corresponded precisely to annotated mouse TSSs (Figure S5). These results show that the putative IRES region of *Hoxa9* encodes a functional promoter.

**Figure 3.**
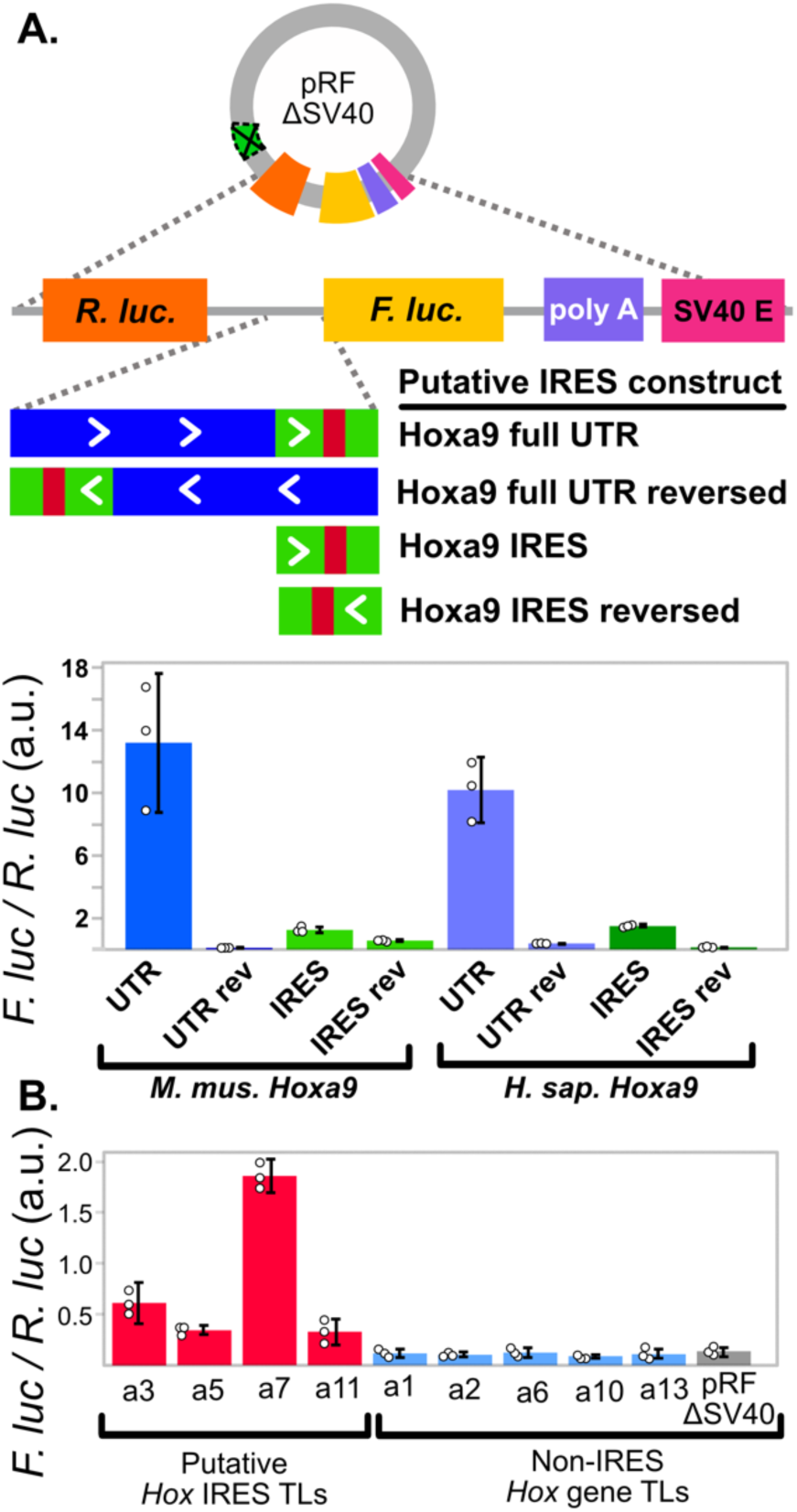
The putative IRES-like domains of *Hoxa9* and other Hox genes encode functional promoters. (A) The putative *Hoxa9* IRES is a promoter. The SV40 promoter was deleted from the pRF bicistronic vector. Putative IRES regions were cloned between Renilla and Firefly luciferase and tested for activity in C3H10T1/2 cells. Bar graphs show the *Fluc* to *Rluc* ratio indicating promoter activity from mouse and human *Hoxa9* regions. The extended UTR and IRES-like regions function as independent promoters in the forward orientation. (B) Putative IRES-like regions from other mouse *Hoxa* genes function as promoters. Annotated transcript leaders from each *Hoxa* gene were tested as in (A). TLs containing putative IRES-like elements drove expression, while non-IRES TLs had background expression levels. Error bars show 95% confidence intervals with n=3.

Based on bicistronic reporter assays, IRES-like activities were previously reported for the annotated TLs from other Hox genes, including *Hoxa3, a4, a5, a7,* and *a11*, while *Hoxa1*, *a2*, *a6*, *a10*, and *a13* did not show activity (3, 17). We next investigated whether the IRES-like regions reported for these genes also have independent promoter activity. Remarkably, all of the previously reported IRES-like TLs we tested (*Hoxa3*, *a5*, *a7*, and *a11*) drove expression of *Fluc* independent of an upstream SV40 promoter, while the non-IRES TLs (*Hoxa1*, *a2*, *a6*, *a10*, and *a13*) had lower *Fluc*/*Rluc* ratios indistinguishable from background noise (Figure 3B). Together, these results suggest the previously reported *Fluc* expression in bicistronic reporter plasmids containing upstream sequences from *Hoxa* genes were due to monocistronic *Fluc* transcripts driven by independent promoters, and not from bonafide IRESes.

We next considered previously identified critical sequences in the *Hoxa9* P4 region. Sequences in the highly conserved 3’ half of this region were previously shown to be required for bicistronic reporter activity and normal skeletal development (3, 4). Specifically, these were sensitive to mutations in the nucleotides underlined “GA**<u>CACGTGAC</u>**<u>”</u>, and similar sequences can be found in other putative *Hoxa* gene IRESes. Using FIMO (32) to search for transcription factor binding sites, we found this sequence matches E-box motifs recognized by at least thirty bHLH transcription factors, including *MYC*/*MAX*, *HES7*, and *ARNT2* (CACGTG) (33, 34), and *USF1*/*USF2*, *ARNTL*, and *TFE3* (CACGTGAC) (34–36). Recent work showed *USF2*binds upstream of human *Hoxa9*, and that co-depletion of *USF1* and *USF2* decreases *Hoxa9* expression in human tissue culture cells (37)(Figure 4A). Similarly, public mouse ChIP-seq data (38–51) show *USF1*, *USF2*, and other bHLH factors bind to the *Hoxa9* E-box, and to E-box regions from other *Hoxa* genes (Figure S7). Remarkably, the *Hoxa9* E-box appears to be universally conserved in vertebrates, as nearly all species evaluated have the *USF1*/*USF2* motif. A CAAT box, considered a core promoter element, is universally conserved adjacent to the E-box, ∼ 60 nucleotides upstream of the major RefTSS annotated TSS (Figures 4B and S8). Notably, the G-rich sequence in the loop region of the mouse *Hoxa9* P4 domain, which has a strong propensity to pair with mouse ES9S through a kissing stem loop interaction (Figure S2), is not similarly conserved in vertebrates. Thus, sequences reported to be essential for mouse *Hoxa9* IRES-like activity encode a deeply conserved transcription factor binding site known to drive expression of human *Hoxa9*.

**Figure 4.**
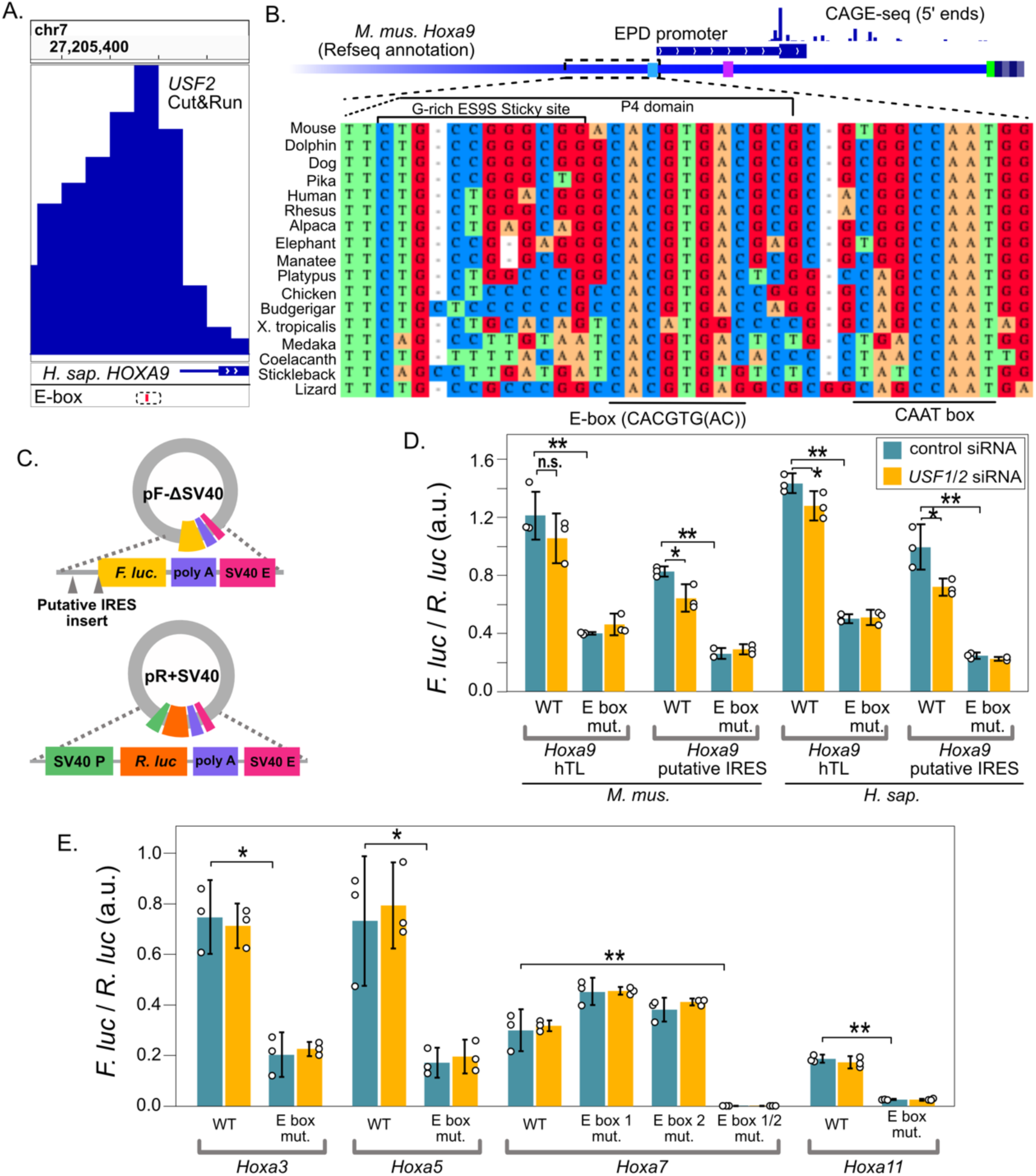
Putative IRES-like elements in *Hoxa* genes contain functional E-boxes recognized by *USF2*. (A) Genome browser view of CUT&RUN sequencing data show the binding location of human *USF2* (37), left. The dashed box shows the location of a hyperconserved E-box motif (also see Figure S7). (B) Sequence alignment from diverse representative vertebrate genomes upstream of mouse *Hoxa9*, including the EPD promoter region. The diagram includes 5’ CAGE-seq data (53). The location of P4 domain sequences and the non-conserved G-rich ES9S interaction site are shown above, while the hyperconserved E-box and CAAT-box are noted below the alignment. The CAAT-box and a TATA-like element are noted on the annotated *Hoxa9* transcript leader in light blue and magenta, respectively (see also Figure S8). (C) The *Fluc* and *Rluc* reporter genes were moved to two independent plasmids to test the functions of E-box elements. (D) Mutation of the E-box motif from mouse and human *Hoxa9* decreased expression in C3H10T1/2 cells. siRNA co-depletion of *USF1* and *USF2* decreased expression of wildtype (Welch’s one-tail t-test, * P < 0.05; ** P < 0.006), but not E-box mutant, reporters. (E) Deletion of E-boxes from putative IRESes of *Hoxa3*, *a5*, *a7*, and *a11* decrease promoter activity. Single E-box mutations show slight increases in *Hoxa7* promoter activity, while the double mutation eliminated promoter function. Error bars show 95% confidence intervals with n = 3.

We tested the hypothesis that these E-box motifs contribute to the promoter activity of Hox gene putative IRES regions. The promoter elements cloned between *Fluc* and *Rluc* in pRF-ΔSV40 plasmids appear to enhance spurious transcription of the upstream *Rluc* gene (15) to various extents (Figures S5-S6; see discussion), which could complicate interpretations of the effect of mutations on *Fluc* expression. Thus, we placed *Rluc* and *Fluc* on separate plasmids to assay the importance of E-box sites in mammalian *Hoxa* genes. Mutating the E-box motifs reduced the promoter activity of mouse *Hoxa3, a5, a7, a11,* and mouse and human *Hoxa9* IRES regions (Figure 4D-E; 1-tail Welch’s t-test P < 0.023). Furthermore, siRNA co-depletion of mouse *USF1* and *USF2* led to a significant reduction in luciferase expression from wildtype mouse and human *Hoxa9* reporters, but not from reporters in which the E-box had been mutated (Figure 4D; 1-tail Welch’s t-test P<0.05). In contrast, *USF1*/*2* co-depletion did not reduce expression from other mouse *Hoxa* putative IRESes (Figure 4E), suggesting their E-boxes may be regulated primarily by other bHLH transcription factors. We conclude that the IRES-like regions of mouse *Hoxa* genes encode functional E-boxes. The function of these sequences as E-boxes explains their necessity for bicistronic reporter expression in previous studies of putative *Hoxa9* IRES activity.

Using the *Hoxa9* gene as a prototypical example of a hyperconserved transcript leader, a recent study identified 589 hTLs in the mouse genome. The authors tested over two hundred of these elements in the bicistronic reporter system and reported ninety (37%) had IRES-like activity (17). Given the misannotation of *Hoxa* gene TLs, we next considered the possibility that these IRES-like hTLs may also be misannotated and encode functional promoters or 3’ splice sites, which both give false-positive results in bicistronic reporter assays. Evaluation of annotated promoter elements (51, 52), TSSs (53), annotated splice sites, and short- and long-read RNA-seq data (54) for TLs reported to have such IRES-like activities revealed the vast majority (85 of 90; 94%) have promoter and / or splicing elements. For example, the *Dedd* gene TL, reported to have the highest IRES-like activity, overlaps two ENCODE promoters and two refTSS sites. ENCODE short- and long-read RNA-seq data support transcription initiation inside the transcript leader, with almost no evidence of full-length hTL expression (Figure 5A). Similarly, approximately one-third of transcripts from *Ptp4a1,* the second strongest IRES-like hTL, initiate within the transcript leader. The IRES-like TL of *Chrdl1* also appears to be misannotated and encodes two EPD promoters with refTSS sites supporting internal transcription initiation (Figure 5A). Other hTLs with reported IRES-like activity overlap known 3’ splice sites (Figure S9A and C). We tested hTLs from four mouse genes that were reported to have IRES activity in C3H/10T1/2 cells (17) - *Chrdl1*, *Cryab*, *Cnot3*, and *Slc25a14*. All showed independent promoter activity in pRF-ΔSV40 (Figure 5B). We also mapped the *Chrdl1*-driven *Fluc* TSS by 5’ RACE and found it matches its annotated EPD promoter and refTSS site (Figure S5). These results suggest that recently reported hTLs with IRES-like activity may also be false-positives, such that their activities in bicistronic reporter assays result from monocistronic transcripts generated from internal promoters and 3’ splice sites rather than bonafide cap-independent translation.

**Figure 5.**
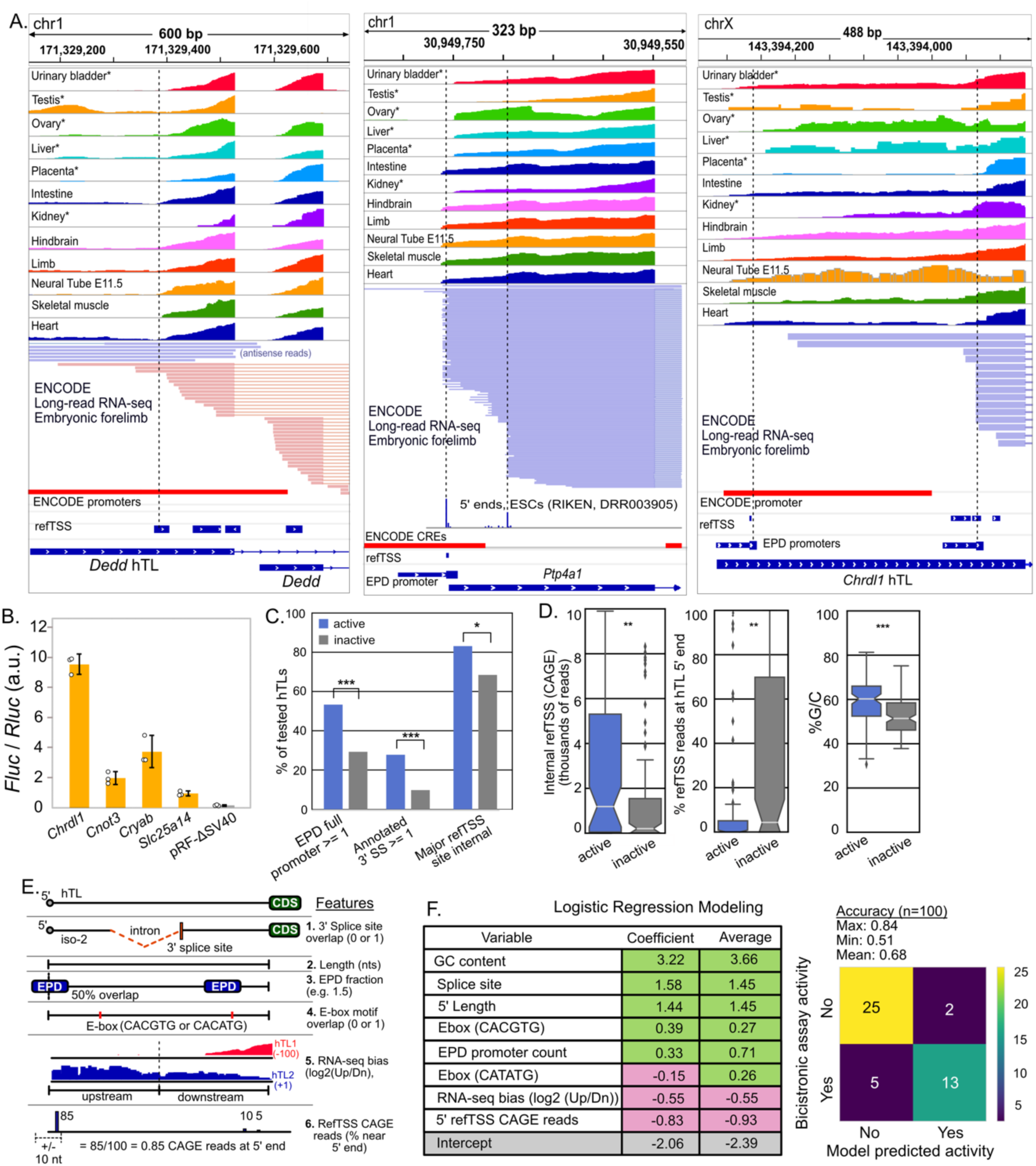
Putative IRES-like hTLs can be explained by promoters and 3’ splice sites due to 5’ UTR annotation errors. (A) Examples of promoter overlap commonly seen in putative IRES-like hTLs. Short-read (upper) and long-read (lower) RNA-seq data show transcription often initiates internally, coinciding with annotated promoters (ENCODE and EPD) and transcription start sites (refTSS). (B) The hTLs from four putative mouse IRESes have promoter activity in pRF-ΔSV40 transfected C3H10T1/2 cells. Error bars show 95% confidence intervals with n = 3. (C) hTLs with putative IRES-like activity are enriched in EPD promoters, 3’ splice sites, and major internal TSS sites (Χ^2^ tests). (D) IRES-active hTLs have significantly more internal CAGE 5’ reads, a lower fraction of TSS reads at annotated 5’ ends and higher G/C content than IRES-inactive hTLs (Wilcoxon Rank-Sum tests). (E) Features for logistic regression modeling. RNA-seq bias is the ratio of reads in upstream and downstream hTL halves across GWIPs-Viz RNA-seq Tables. RefTSS CAGE reads are the percentage of 5’ end reads mapped near the annotated transcription start site (data from 28). (F) Logistic regression modeling of IRES-like and non-IRES hTLs. Features associated with internal promoters (GC content, EPD promoter count fraction, E-boxes) and splice sites are positively correlated with bicistronic reporter expression, while features associated with full length TLs (CAGE reads at annotated 5’ ends and RNA-seq 5’ end bias) are negatively correlated with bicistronic reporter activity. 100 models were generated, with an average accuracy of 68%. *, **, and *** denote P < 0.05, 0.01, and 0.001, respectively.

We next systematically evaluated the 589 previously reported hTLs for potential misannotation due to overlapping promoters, enhancers, TSSs, 3’ splice sites, and protein coding sequences (CDSs), all of which could contribute to high conservation rates unrelated to translational control. Using combined promoter/enhancer sets, we find 93% of hTLs overlap ENCODE (509) and/or EPD (463) promoters (Figure S9B). Of those, 221 hTLs contain full-length EPD and 100 contain full-length ENCODE promoters. To further evaluate the accuracy of mouse TL annotations, we examined refTSS annotations and their underlying quantitative 5’-CAGE high-throughput sequencing data (53). Of the hTLs with public 5’ CAGE data, only 40% have an annotated refTSS site within 10 nucleotides of their annotated 5’ ends. Furthermore, 78% of all TSS-containing hTLs had the strongest CAGE peak within the hTL rather than near the 5’ end. The complexity of the mouse transcriptome further complicates conclusions about TL conservation, as 17% of hTLs overlap annotated 3’ splice sites and 43% of hTLs overlap annotated CDSs from alternative transcript isoforms (Figure S9C). Indeed, one-third of hTLs had CDS overlap covering at least 25% of their length and one-fifth were at least 50% overlapped by annotated CDSs (Figure S9D). These data show that many recently reported hTLs are misannotated such that their high conservation rates may reflect evolutionary pressure to maintain promoters, enhancers, splice sites, and protein coding sequences.

Our luciferase reporter assays show clear evidence that previously reported IRES-like elements result from transcriptional promoter activity. To further evaluate the potential for such false positives in recently reported hTLs, we compared the frequencies of promoters and splice sites, and the distribution of 5’ CAGE reads in hTLs that were reported to be “active” (90) and “non-active” (133) in the bicistronic assay. Active hTLs are 1.7-fold more likely to overlap at least one complete EPD promoter than “non-active” hTLs (53% vs 29%; Chi-squared test P = 0.0002107, Figure 5C). Similarly, annotated 3’ splice sites were 2.8-fold enriched in active, as compared to non-active, hTLs (28% vs 10% Chi-squared test P = 0.000451, Figure 5C). Furthermore, ∼83% of active TLs have their strongest CAGE peak within the TL compared to 68% for the non-active TLs (Chi-squared test P = 0.02009, Figure 5C). Finally, active hTLs have drastically more CAGE reads at internal refTSS sites (P = 0.00144), a lower fraction of CAGE reads near their annotated 5’ ends (P = 0.00173), and higher GC content (P = 2 x 10^-6), which is characteristic of promoter elements (Figure 5D) (55). The enrichment of promoters, splice sites, and internal 5’ CAGE reads in bicistronic “active” hTLs suggests that these elements generally drive bicistronic reporter expression through the creation of monocistronic *Fluc* transcripts, rather than by cap-independent IRES-like activities.

If internal promoters and splice sites are responsible for the reported IRES-like activities in hTLs, we reasoned that such features could be used to predict their activities. To test this, we used logistic regression with these features to model their activity in the bicistronic reporter assay (Figure 5E-F; see Methods). Strikingly, this approach generated models that were, on average, 68% accurate at predicting IRES-like activity (maximum accuracy 84%). Features associated with promoters (GC content, EPD promoter counts, E-boxes) and 3’ splice sites were positively correlated with bicistronic active hTLs, while those reflecting high levels of full length hTL transcription (5’ refTSS fraction and 5’ RNA-seq bias) were associated with inactive hTLs. The ability to accurately predict bicistronic assay activity from these genomic features strongly supports the conclusion that such activities are false-positives, inconsistent with their putative functions as IRES-like elements driving cap-independent translation.

## Discussion

The vast majority of mRNAs are believed to undergo cap-dependent translation in rapidly dividing cells, while cap-independent mechanisms, including IRESes, are used primarily during cell stress (2). Over the last decade, multiple studies have coalesced on an intriguing model proposing key developmental genes are regulated by cap-independent translation driven by IRES-like sequences and structures in hyperconserved 5’ transcript leaders (hTLs). However, this model was founded on bicistronic reporter assays, which are subject to common false-positive results due to cryptic promoter and splicing activities. Here, we examined previously reported hTLs with putative IRES-like activities in *Hoxa9* and other genes. We found many putative IRES regions are not transcribed at biologically relevant levels. In addition, the putative IRESes that are transcribed are enriched in internal promoters and / or 3’ splice sites known to cause false-positives in bicistronic reporter assays. Consistent with this, a concurrent independent study found much shorter 5’ UTRs in *Hox* genes and showed the putative *Hoxa9* IRES has promoter activity in tissue culture cells (56). Furthermore, we successfully predicted putative IRES-like activity using known annotated promoters, 3’ splice sites, CAGE-seq 5’ end data and public RNA-seq data. Finally, we find promoters, splice sites, enhancers, and even protein coding sequences overlap hTLs, which may explain their sequence conservation. Our results provide conventional explanations for unconventional results from previous studies, requiring a reevaluation of the proposal that these TLs drive cap-independent translation.

We found the putative *Hoxa9* IRES has only trace evidence of expression in mouse RNA-seq data and instead encodes a functional promoter. Consistent with this, our R-scape analysis, made possible by recently published mammalian genome sequences (21), indicates the proposed RNA structure of the *Hoxa9* IRES (3) is not constrained by evolution. Instead, the P4 region encodes a hyperconserved E-box motif recognized by *USF1*/*USF2* whose mutation drastically decreases promoter activity. Three recently reported mutations to this E-box, M3, M5, and M8 (4), modified, 1, 4, and 3 nucleotides, respectively, and decreased expression in the bicistronic reporter assay, as expected for loss of a transcription factor binding site. Notably, the effects of these mutations corresponded to the number of nucleotides modified, as M3 had a smaller effect than M5 and M8. However, the M5 mutation also appeared to shift *Hoxa9* mRNA from the polysome towards the monosome (4). While this may seem to support IRES-like elements, it can also be explained by promoter activity. Mutating the E-box likely decreases the production of the natural 83-nt TL isoform, such that spurious longer transcripts, unspliced transcripts, and *Hoxa9*/*a10* fusion transcripts (Figure 2) make up a larger fraction of *Hoxa9* mRNA in polysome gradients. These longer transcripts include up to 14 uORFs and would not be translatable but would be detected using the RT-qPCR primers from these studies, which are not specific to spliced, mature *Hoxa9* mRNA (Figure 2). Similar reasoning could explain how deletion of the putative IRES region appeared to alter *Hoxa9* polysome association in mouse embryos based on RT-qPCR (3). Thus, the intrinsic promoter activity we observed in *Hoxa9* genes provides a conventional explanation for the effects of these mutations, especially since the putative IRES is present in only ∼1% of mRNA transcripts *in vivo* (Figure 2) (56).

A recent study also proposed the *Hoxa9* P4 stem-loop recruits translation preinitiation complexes (PICs) through interactions with ribosomal expansion segment ES9S (4). However, these assays were performed at 4°C and may not be physiologically relevant. Indeed, the P4 domain and ES9S have the potential to form kissing loops with nine G-C and one G-U base pairs, which appears consistent with a published cryo-EM structure (Figure S2)(4). This interaction has a predicted free energy of -15.94 kcal/mol (RNAcofold)(57) and would thus be very stable under cryo-EM and affinity purification conditions. Notably, neither the P4 structure nor the G-rich stretch of *Hoxa9* were required for IRES-like activity (4), and neither are evolutionarily conserved (Figures 1 and 4). Because the putative IRES structure is not conserved, and is rarely, if ever, expressed as a transcript leader (Figures 2, S3 & S4,(56)), our results contradict the notion that mammalian ES9S recruits PICs to *Hoxa9* for cap-independent translation. A transcriptome-wide screen reported that ES9S similarly binds to mouse mRNA fragments with G-rich motifs, several of which were reported to have IRES-like activity using the bicistronic reporter system (5). However, these putative IRESes often overlap promoter elements (Figure S10) and no controls were performed to test for false-positives from monocistronic *F. luc.* Consequently, we propose that the interactions observed *in vitro* between the ES9S and G-rich mRNA are coincidental associations stabilized by low temperature.

The TLs of *Hoxa3*, *a4*, *a5*, *a7*, and *a11* were also previously reported to have IRES activity based on bicistronic assays (3, 17). As with *Hoxa9*, many of the other previously reported *Hoxa* gene IRES-like TLs appear to be misannotated. For example, Xue et al. defined a 1,106 nt TL for *Hoxa4* using 5’ RACE (3). However, the contemporaneous transcript annotation indicated a 15 nt leader, which is supported by RNA-seq data (Figure S11). Similarly, the 1,168 nt and 496 nt IRES-like TLs from mouse *Hoxa7* and *a11* appear to be only ∼112 nt and ∼90 nt long, respectively (Figure S10). Overall, our results suggest *Hoxa* mRNAs have shorter TLs translated via cap-dependent translation. Since it is much more efficient in developing embryos than cap-independent translation (58), cap-dependent translation would help ensure robust timely expression of these key developmental regulators.

Consistent with their misannotation, all the putative *Hoxa* IRESes we tested (*a3*, *a5*, *a7*, and *a11*) showed independent promoter activities, while non-IRES *Hoxa* TLs did not. Strikingly, sequences previously shown to be sufficient for putative *Hoxa3, a4, a5,* and *a11* IRES activities (3) overlap annotated promoters and TSSs (Figure S11). Moreover, conserved E-boxes were found in all the *Hoxa* TLs with putative IRESes (Figure S11), but not in non-IRES TLs. Mutating these E-box sites decreased the strength of *Hoxa* promoter activities. We showed depletion of *USF1* and *USF2* decreased expression from the mouse and human *Hoxa9* reporters. However, mutating *Hoxa9* E-boxes had a stronger effect (Figure 4), suggesting other bHLH transcription factors may also promote expression from these binding sites. Along those lines, other *Hoxa* reporters were not affected by *USF1*/*USF2* depletion, suggesting they may instead be regulated by other bHLH factors that recognize the same core motif (CACGTG) (33, 36). Notably, public mouse ChIP-seq data show *USF1*, *USF2*, *MYC*, *MAX, TFE3*, *ARNTL*, *BHLHE40*, and *BHLHE41* bind to *Hoxa* gene E-boxes (Figure S7) (38–42, 44–51, 59–66). Several other E-box recognizing transcription factors, including *TCF15*, *HES1*, *HES7*, *MESP2*, and *MSGN1*, have been implicated in somite formation (67–72), and may also regulate *Hoxa* genes. More studies are needed to investigate the functions of conserved E-boxes in regulating *Hox* gene transcription during development.

Translational control of *Hox* genes was first suggested by a report that their translation was reduced in mouse embryos hemizygous for *RPL38* (*Ts/+*) (58). However, the data presented in that study do *not* actually show a decrease in Hox mRNA translation, typically seen as a shift from polysome to monosome sucrose gradient fractions. Instead, *Hox* mRNA were substantially *reduced in both polysomes and monosomes* in Ts/+ embryos, though the data were presented in separate figures (Figures 3 and 6 from (58)). Furthermore, only a slight increase was observed in non-translating fractions (58)). Although the authors reported *Hox* gene mRNA levels were not decreased in *Ts/+* mutant embryos, the underlying RT-qPCR results had such high variance that even considerable changes in mRNA levels would be undetectable, and the *Hoxa9* primers used would not distinguish between mature mRNA, unspliced pre-mRNA, or fusion transcripts (Figure 2). Even with optimal primers, qPCR has several limitations in estimating mRNA levels (73–75). Additionally, a recent ribosome profiling study in HEK293 cells reported that depletion of *RPL38* decreased the translation efficiency of many genes that promote WNT signaling and *Hox* gene transcription (76). Future ribosome profiling studies from wildtype and Ts/+ embryos are needed to determine whether *RPL38* hemizygosity actually disrupts translation of *Hox* genes, their upstream transcriptional regulators, or both.

Our results also do not support the recently reported catalog of 589 hTLs in other mouse genes (17), ninety of which have putative IRESes based on bicistronic reporter assays. We showed these hTLs frequently overlap annotated promoters, enhancers, 3’ splice sites, and even protein coding sequences, providing conventional explanations for their unusually high conservation rates. Furthermore, we tested four putative IRES regions from these hTLs and found all encoded promoters. Indeed, the ninety IRES-like hTLs often show internal transcription initiation in public RNA-seq from ENCODE and 5’ CAGE-seq from RIKEN, and are particularly enriched in annotated promoters and splice sites, compared to non-IRES hTLs. We also used these features to build a model predicting bicistronic reporter activity. Notably, this model showed GC content and length were positive predictors of bicistronic activity – features that might appear consistent with structured IRESes. However, a high-throughput bicistronic IRES screen with controls to reduce promoter and splicing artifacts previously showed GC content was lower in active IRES elements (16) and high GC content is a known hallmark of promoter regions (55). Taken together, we propose that these hTLs are incorrect due to transcriptome annotation errors and promoter and splicing activities in bicistronic reporter assays. However, the concept of hyperconserved elements in 5’ TLs is still intriguing and deserves more careful study to identify genuine hTLs and investigate their functional elements.

It is well known that bicistronic reporter assays are subject to false positive results due to cryptic promoters and splice sites. Many control experiments have been devised to account for this. These include RNAi treatment to identify monocistronic *Fluc* transcripts, RT-PCR screening for cryptic splicing, and deletion of the SV40 promoter upstream of *Rluc* to account for independent promoter activities (7, 8, 20). Notably, a previous study of Hox gene IRES activity used siRNA targeting *Rluc* as a control for monocistronic transcripts. If only bicistronic transcripts were present, this treatment should deplete both *Rluc* and *Fluc* mRNA. Although siRNA treatment nearly eliminated *Rluc* mRNA, ∼30% of *Fluc* mRNA remained, consistent with monocistronic *Fluc* expression driven by promoter activities from the *Hoxa3*, *a4*, *a5*, *a9*, and *a11* IRES-like regions (3). Our results further support such monocistronic transcripts, as the putative IRES-like *Hoxa* TLs we tested had independent promoter activity, while non-IRES *Hoxa* TLs did not.

Previous work showed that the pRF plasmid has two cryptic promoters upstream of *Fluc* and *Rluc* which generate a variety of cryptic spliced products (Figure S1) (15). Using 5’ RACE, we identified *Rluc* transcripts containing multiple uORFs from one of these cryptic promoters, in the pMB1 origin of replication (Figure S5). We suspect the expression of *Rluc* by putative IRES-test sequences (Figure S6) may reflect induction of other spurious transcripts, perhaps initiating at the other known cryptic promoter in the f1 origin of replication (15)(Figure S1). Regardless of the mechanism, the existence of these spurious transcripts further undermines comparisons of *Fluc*/*Rluc* protein and RNA ratios, previously used to discount cryptic promoters and splicing of putative IRES-like hTLs (17). These issues have been previously noted(7, 8, 15), with arguments specifically against using RT-qPCR for bicistronic assays because it is unclear which transcripts are amplified in such assays (20).

Considering the cryptic promoters and splicing events associated with the pRF plasmid (15), IRES studies using this vector require rigorous controls (*Rluc*. RNAi, promoter deletion, *Fluc* 5’ RACE) to eliminate the possibility of monocistronic transcripts from each test IRES sequence. However, it may be preferable to completely forego use of the bicistronic reporter. Because putative IRES sequences could alter the expression of spurious *Rluc* transcripts containing various numbers of uORFs (15)(Figure S1, S5), which likely have variable mRNA stability and translation efficiency, *Rluc* mRNA and protein levels may also not be reliable as internal controls in the bicistronic reporter. Given these complications, we instead advocate testing IRES activity by comparing reporter expression from directly transfected m7G- and A-capped linear transcripts, using circRNA reporters, or both (7, 8, 77, 78).

Our results underscore the importance of accurate transcript annotations for defining and studying TLs. The incorrect, extended *Hoxa9* TL can be traced to experiments that used reverse transcription “primer walking” to find the most upstream 5’ end (79). Unfortunately, this appears to have also amplified introns from *Hoxa9* fusion transcripts. Indeed, the region upstream of this misannotated extended TL is extremely G-rich, such that G quadruplexes may have halted reverse transcription. As recently noted (56), the annotated mouse *Hoxa9* transcripts are 600-800 nt longer than expected given northern blots in prior work (79, 80), further indicating their misannotation. However, such annotation errors are common, as many extended TLs from other genes also appear to be incorrect (e.g., *Hoxa4*, *Hoxa7*, etc.). In other cases, transcription initiates at multiple sites within annotated TLs (e.g., *Ptp4a1*, *Chrdl1, Dedd*). Astonishingly, even the TL of mouse *Actb* (beta actin) appears to be misannotated in RefGene, initiating with a TATA box and including a promoter (Figure S12). This error may explain why its TL showed apparent IRES-like activity when fused to the *Hoxa9* P4 domain (4). Similar errors may underlie additional putative IRESes from mRNAs that bind to ES9S in vitro (5), as most of these also include annotated promoter regions or other evidence of internal transcription initiation (Figure S9), Added to this is the general complexity of mammalian transcriptomes, in which TLs often include promoters, introns and 3’ splice sites. Together, these issues make accurate mammalian TL annotation particularly challenging, and complicate the study of TL functional elements and conservation. Ongoing efforts to sequence full length transcripts (81, 82), integrated with annotated promoters and TSSs, should eventually resolve such issues and greatly aid the study of TL functions in mammals.

## Materials and Methods

### Luciferase vector cloning

The pRF+423Dux4 plasmid (Addgene #21625) contains Renilla Luciferase (R-luciferase) under the control of a SV40 promoter. Firefly luciferase, downstream of R-luciferase, is transcribed under the control of the same SV40 promoter and is preceded by a putative upstream IRES (Figure S1). The pRF+423Dux 4 vector was sequenced using primers that anneal to the pGL3 vector (Promega, see primers Table S2). The Dux4 IRES site was deleted from pRF+423Dux4 using PCR primers that flank the IRES region (PRF423DUX4-ATW F and R, Table S2). The primers also incorporated a BglII site after the start codon of F-luciferase, with an upstream HindIII site. The PCR product was phosphatased and circularized by ligation to create the vector pRF-ΔIRES. To delete the SV40 promoter, add an EcoRI site, and remove an additional BglII site, pRF-ΔIRES was used as a template for a second PCR, using the primers SV40D-EcoRI and SV40D-XhoI-R (Table S2). The resulting PCR product was phosphatased and circularized by ligation to create the vector pRF-ΔSV40. Both pRF-ΔIRES and pRF-ΔSV40 were verified by Sanger sequencing and tested for luciferase activity in C3H/10T1/2 mouse embryonic fibroblast cells (MEF) obtained from ATCC. A R-luciferase only vector was constructed by removing F-luciferase from pRF-ΔIRES by XbaI digestion. The XbaI cut vector was gel purified and circularized by ligation, resulting in the vector pR+SV40. The pR+SV40 vector was Sanger sequenced and its luciferase activity was verified in MEF cells.

### Putative 5’UTR cloning

Putative 5’UTR sequences were obtained as double stranded DNA fragments from Twist Biosciences and Genewiz (Table S2). The DNA fragments were PCR amplified, digested with HindIII and BglII, and cloned into the pRF-ΔSV40 vector at the HindIII and BglII sites upstream of F-luciferase. Due to limitations in DNA synthesis, five additional As were added by site-directed mutagenesis using MMHOXA11-ATW forward and reverse primers (Table S2) to finish the *Hoxa11* construct. Additional sequences in the *Hoxa3* hTL were removed by PCR using primers HOXA3-SATW forward and reverse primers (Table S2). Site-directed mutagenesis was also performed on Hoxa3, a5, a7, a9, and a11 constructs to mutate E box sites to CACTAT. Some inserts affected both F-luciferase and R-luciferase (Figure S6), For more precise ratiometric measurements, the R-luciferase gene was removed from wildtype and E-box mutant *Hoxa3*, *a5*, *a7*, *a11*, and *a9* constructs by EcoRI and HindIII digestion, end polishing with DNA polymerase I large fragment (Klenow), and re-ligation (pF-ΔSV40; Figure 4C). All constructs were Sanger sequenced (Table S2), and transfection grade plasmid DNA was purified using a Qiagen Plasmid Mini column according to the manufacturer’s instructions.

### Luciferase Assays

In 96-well tissue culture plates, 2 x 10^3^ mouse embryonic fibroblast (C3H/10T1/2 clone 8, ATTC) cells seeded in 100 μl DMEM supplemented with 10% FBS per well. Cells were allowed to adhere and grow for 24 hours at 37 °C. In 10 μl of Opti-MEM™. 100 ng of construct was mixed with 0.4 μl of ViaFect™(Promega) and incubated for 12 minutes at room temperature. The transfection mixture was added dropwise to the cells and the cells were incubated at 37 °C for 24 hours. F-luciferase and R-luciferase expression was assayed in a TECAN Spark plate reader using the Dual-Glo® Luciferase Assay System (Promega) according to the manufacturer’s instructions. Both F-luciferase and R-luciferase were measured for 10 seconds per well.

### *USF1* and *USF2* siRNA knockdown

In a 96 well tissue culture plate, 1 x 10^3^ mouse embryonic fibroblast (C3H/10T1/2 clone 8, ATTC) cells seeded in 100 μl DMEM supplemented with 10% FBS per well. Cells were allowed to adhere and grow for 24 hours at 37 °C. In 10 μl of Opti-MEM™, 1 pmol of siRNAs (scrambled control or USF1/2 siRNA; Santa Cruz Biotechnologies) were mixed with 0.3 μl of Lipofectamine™ 3000 transfection agent and incubated at room temperature for 15 minutes. The transfection mixture was added dropwise to the cells so that the final concentration of each siRNA was 10 nM. The cells were incubated at 37 °C for 24 hours. For each well, 20 ng of pR+SV40 vector (Renilla only) and 80 ng of a Hox gene construct were mixed with 10 μl Opti-MEM™ and 0.5 μl of ViaFect™ and incubated for 15 minutes at room temperature. The mixture was added dropwise to the cells and the cells were incubated at 37 °C for 24 hours. F-luciferase and R-luciferase were assayed as described above (Luciferase Assay).

### Validation of *USF1* and *USF2* siRNA knockdown

In a 6 well tissue culture plate, 3 x 10^4^ C3H/10T1/2 cells seeded in 2 ml DMEM supplemented with 10% FBS per well. Cells were allowed to adhere and grow for 24 hours at 37 °C. In 250μl of Opti-MEM™, 20 pmol of siRNAs (either scrambled control or USF1/USF2 siRNA) were mixed with 7.5 μl of Lipofectamine™ 3000 transfection agent and incubated at room temperature for 15 minutes. The transfection mixture was added dropwise to the cells so that the final concentration of each siRNA was 10 nM. The cells were incubated at 37 °C for 48 hours. The media were removed, and total RNA was extracted using TRIzol™ (Invitrogen) following the manufacturer’s instructions.. The total RNA was twice treated with TURBO™ DNase (Invitrogen™) and purified over an RNA Clean and Concentrator -5 column (Zymo Research) after each DNase treatment. RT-qPCR was performed in 50μl reactions using the SuperScript™ III Platinum™ SYBR™ Green One-Step RT-qPCR kit (Invitrogen™) with 200 ng of total RNA as template. Cycling and reaction conditions were followed according to the manufacturer’s instructions (see Table S2 for primer sequences). Three biological replicates were performed for the knockdown and scrambled control. Three technical replicates were performed for each gene, along with three technical replicates of no template controls. No amplification was detected for the no template controls. Relative gene expression of USF1 and USF2 were compared to GAPDH using the ΔΔ-Ct method (Figure S13).

### Logistic Regression

We used active and non-active hTLs provided by Gun Woo Byeon and Maria Barna (17). After removing records from TLs that were not previously classified as hyperconserved, the Table included 133 nonactive hTLs and 90 active hTLs. We compiled a list of several categorical and numerical sequence features that could contribute to bicistronic activity (e.g. GC content, CAGE data, ebox motifs; Figure 5; Table S3). Transcript leaders lacking sufficient data for refTSS calculations (5’ RefTSS CAGE reads) were assigned a mean imputation filler value. To perform classification of active versus non-active transcript leaders, we used LogisticRegressionCV from scikit learn (sklearn.linear_model.LogisticRegressionCV) with the default solver=lbfgs, Cs = 10, intercept=True, and cv = 10 parameters. All numerical features were normalized using sklearn.preprocessing.MinMaxScaler. 100 separate models were individually trained on random samples of 80% of the data and tested on the remaining 20% of the data.

### ENCODE RNA-Seq Data

From the ENCODE database, we used polyA plus RNA-seq data from *Mus. musculus* and *Homo sapiens* tissues that showed evidence of transcription for *Hoxa9*, assessed by visual examination. For the tissues containing multiple bigwig files, we merged the reads to create a new complete bigwig. The files containing “negative strand signal” data were used, because *Hoxa9* is on the negative strand. If no negative specific file was given, then the “all reads” signal was used. For positive strand gene examples, the “positive strand signal” files were used instead. The accession numbers used are included in Table S4.

### CAGE data (RefTSS)

CAGE-seq data were downloaded using SRA Run Selector from NCBI from SRA study number DRP000949 (BioProject PRJDB1980). In this study CAGE reads were obtained from Human and Mouse transcripts to define transcription start sites (TSSs). For our study, we used the Mus musculus data from runs DRR003905 (experiment DRX003141) and DRR003906 (experiment DRX003142). The data were from induced pluripotent stem cells (iPCSs) and embryonic stem cells (ESs) respectively. Reads were processed using fastq-dump followed by cutadapt. The processed data was aligned to the mouse genome using STAR. Reads were summed and assigned to annotated refTSS peaks via bedtools intersect to define refTSS strength. Files used are listed in Table S4.GWIPS RNA-seq data was retrieved from GWIPS table browser (83) from Mouse (mm10) using the global aggregate setting. Data was compiled from 26 files (listed below). Bedgraphs were combined using bedtools unionbedg.

Alvarez Dominguez 2017, Atger 2015, Atlasi 2020, Blanco 2016, Castaneda 2014, Castela Szekely 2017, Cho 2015, Eichhorn 2014, Fradejas-Villar 2016, Freimer 2018, Gonzalez 2014, Guo 2010, Hornstein 2016, Howard 2013, Hurt 2013, Ingolia 2011, Janich 2015, Laguesse 2015, Rapino 2018, Reid 2014, Reid 2016, Reid 2017, Sendoel 2017, Simsek 2017, Thoreen 2012, You 2015

### Infernal & Rscape

The sequence for the predicted mouse *Hoxa9* IRES from (3) was used. Using the latest Zoonomia Cactus alignment file, we mapped the Mouse coordinates (chr6:52226238-52226413) to 208 vertebrate species via halLiftover (84). An additional 23 vertebrate sequences were extracted from the UCSC database for a total of 231 species. The sequences and results are in Table S1. The putative IRES structure from (3) was converted into dot-bracket notation and used to generate a Stockholm format file containing the 231 sequences, the conserved structure, and the *Mus musculus* sequence as reference.

The Infernal package was used to build a covariance model and prune the sequence alignment. Using default parameters for cmbuild and cmcalibrate, 25 close species were used to build and calibrate the initial covariance model. Using cmsearch, target sequences (230 sequences) with appropriate E-values (default) for covariation testing were kept (medaka not significant, filtered out). A new Stockholm file was generated from the remaining sequences. The resulting file was used as input to R-scape using default parameters with various E-value thresholds (0.005, 0.01, 0.1, 1, 10, 20, 30). The IRES data were tested seven times with the varying E-values.

### 5’ RACE from pRF reporter plasmids

3.2 x 10^5^ C3H/10T1/2 cells (clone 8, ATTC) were seeded in 10 ml DMEM (10% FBS) in 10 cm dishes and grown for 24 hours at 37 °C. 17 μg plasmid, 43 μl of Lipofectamine 3000™, and 34 μl of P3000 reagent (ThermoFisher) were combined in 1 ml of Opti-MEM™, incubated for 15 minutes at room temperature, and added dropwise to the cells. The cells were incubated at 37 °C for 24 hours and harvested in 2 ml of TRIzol™ (Invitrogen™). RNA was extracted following the manufacturer’s instructions. The RNA was pelleted by centrifugation at 20,000 G for 30 minutes at 4 °C. The pellet was washed with 70% ethanol and resuspended in 200 μl of nuclease free water. 45 μg of total RNA was twice treated with TURBO™ DNase (Invitrogen™) and purified over an RNA Clean and Concentrator -25 column (Zymo Research). The RNA was eluted in 200 μl of nuclease free water, and mRNA was selected using 75 μl of Oligo d(T)25 magnetic beads (New England Biolabs) according to the manufacturer’s instructions. Poly-A mRNA was eluted in 36 μl of nuclease free water and 12 μl was reverse transcribed in a 30 μl reaction using the Template Switching RT Enzyme Mix (New England Biolabs) according to the manufacturer’s instructions. Primers LUC-RT-R2, and Rluc-RT-R (Table S2) were used for F- and R-luciferase, respectively, and a mix of both primers were used in no-RT controls. The template switching oligo (TSO-Eno2, Table S2) adds a forward primer site for PCR. cDNA were purified with AMPure® XP magnetic PCR purification beads (Beckman Coulter), eluted in 10 μl of nuclease free water, and PCR amplified for 35 cycles using the primers, ENO2LIBF1 and LUC-R for F-luciferase and ENO2LIBF1 and R-LUC-int-R for R-luciferase, in a 25 μl reaction with Phusion® High-Fidelity DNA Polymerase (New England Biolabs) in high GC content buffer with DMSO according to the manufacturer’s instructions. The PCR products were visualized on a TapeStation (Agilent). PCR products were electrophoresed on a 2% agarose gel, and visible bands were excised, gel extracted, and cloned using the Zero Blunt™ TOPO™ PCR cloning kit (Invitrogen™) according to the manufacturer’s instructions.

## Acknowledgments

We would like to thank Cassia Williams-Rogers for assistance in computational data analysis and John Woolford for helpful discussions and comments on the manuscript. We thank Gun Woo Byeon and Maria Barna for providing a table of hTL bicistronic assay data previously summarized in (17). This study was supported by NIH R01 GM121895 to CJM.

---

**Figure S1.**
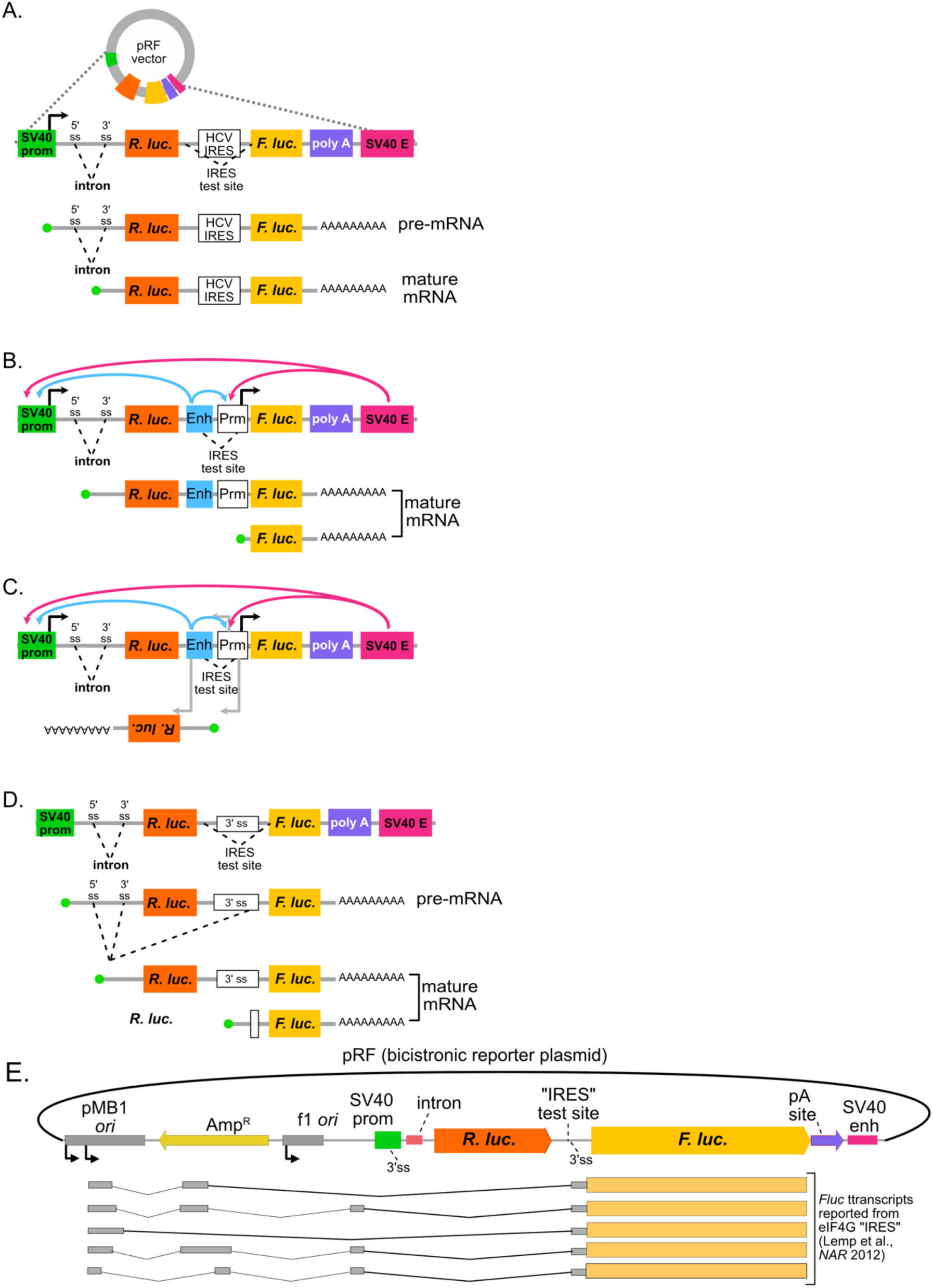
The pRF bicistronic reporter is subject to cryptic promoter and splicing activity. (A) Design of the pRF vector and expected mRNA products. Transcription is driven by the SV40 promoter (green), increased by an included SV40 enhancer (magenta). An intron is positioned upstream of Renilla luciferase (*Rluc*) to increase expression. An IRES test site between *Rluc* and Firefly luciferase (*Fluc*) ORFs is shown with the Hepatitis C Virus IRES (HCV). Transcription and splicing are expected to generate homogenous bicistronic transcripts. **(B)** Enhancer and promoter sequences placed in the IRES test site can generate monocistronic *Fluc* transcripts that give false-positive “IRES” signals. The “IRES” enhancer sequence (blue) can alter the transcription of both bicistronic and monocistronic transcripts. **(C)** The bidirectional transcription from promoters and enhancers may produce antisense *Rluc* transcripts, which would complicate qRT-PCR normalization attempts. (D) 3’ splice sites (both strong and cryptic) in the IRES test site also create false-positive *Fluc* expression from monocistronic transcripts. **(E)** Lemp et al., (*NAR* 2012) showed the pMB1 and f1 replication origins have promoters that drive aberrant *Fluc* products via cryptic splicing. Arrows indicate transcription start sites. Aberrant transcripts from the eIF4G test “IRES” are shown. Cellular transcripts were also *trans*-spliced to *Fluc* in transfected tissue culture cells. Similar transcripts can affect *Rluc* (see Figure S5) and also leave additional *Rluc* RNA in spliced lariat introns. This likely undermines *Rluc* / *Fluc* qRT-PCR assays.

**Figure S2.**
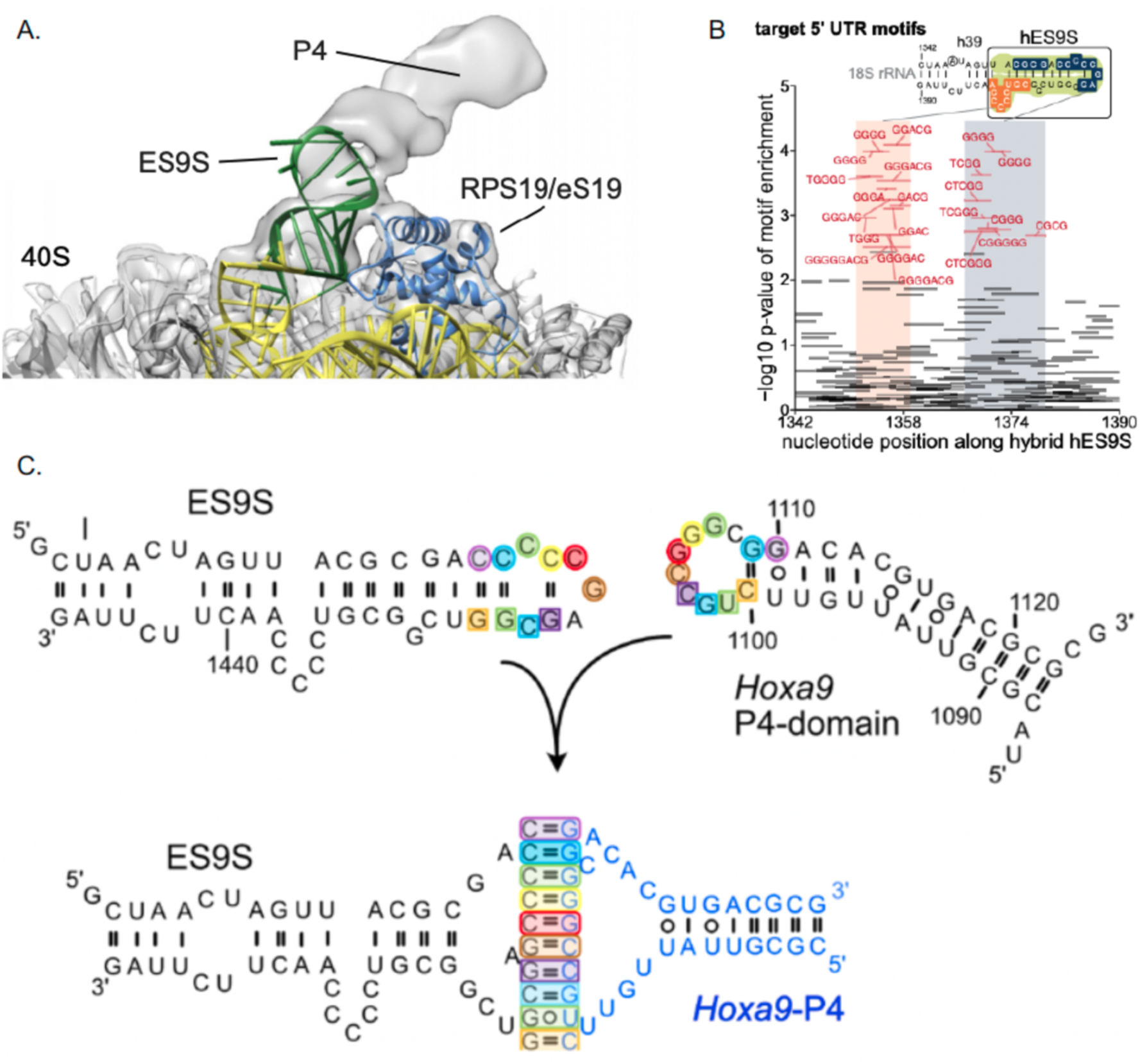
Models of interactions between ES9S, *Hoxa9* “P4” and mouse mRNA fragments. (A) The Cryo=EM interaction model published by Leppek et al., 2020. An A-form helix labelled “P4” appears oriented toward the C-rich loop of ES9S (green). (B) Leppek et al., 2021 figure panel 5a, showing G-rich motifs enriched in fragmented mouse mRNAs tthat bound to ES9S in vitro. The ES9S sequence was depicted above, with C-rich regions highlighted, as Leppek et al proposed these might form complementary Watson-crick pairs. (C) Mouse ES9S and putative *Hoxa9* IRES P4-domain have complementary sequences that could support a kissing stem loop interaction. Proposed structures of individual RNAs are depicted as reported from Leppek et al., 2020 and Xue et al., 2015 (above). Nucleotides that have the propensity to pair are shaded in matched colored ovals and squares. A G-rich segment in the putative *Hoxa9* IRES P4-domain is complementary to a C-rich segment in ES9S, with further potential pairing between additional adjacent nucleotides. Note the similarity to the interaction proposed by Leppek et al. 2021 (panel B) for ES9S binding to G-rich motifs in other mouse mRNAs (except *Hoxa9*).

**Figure S3.**
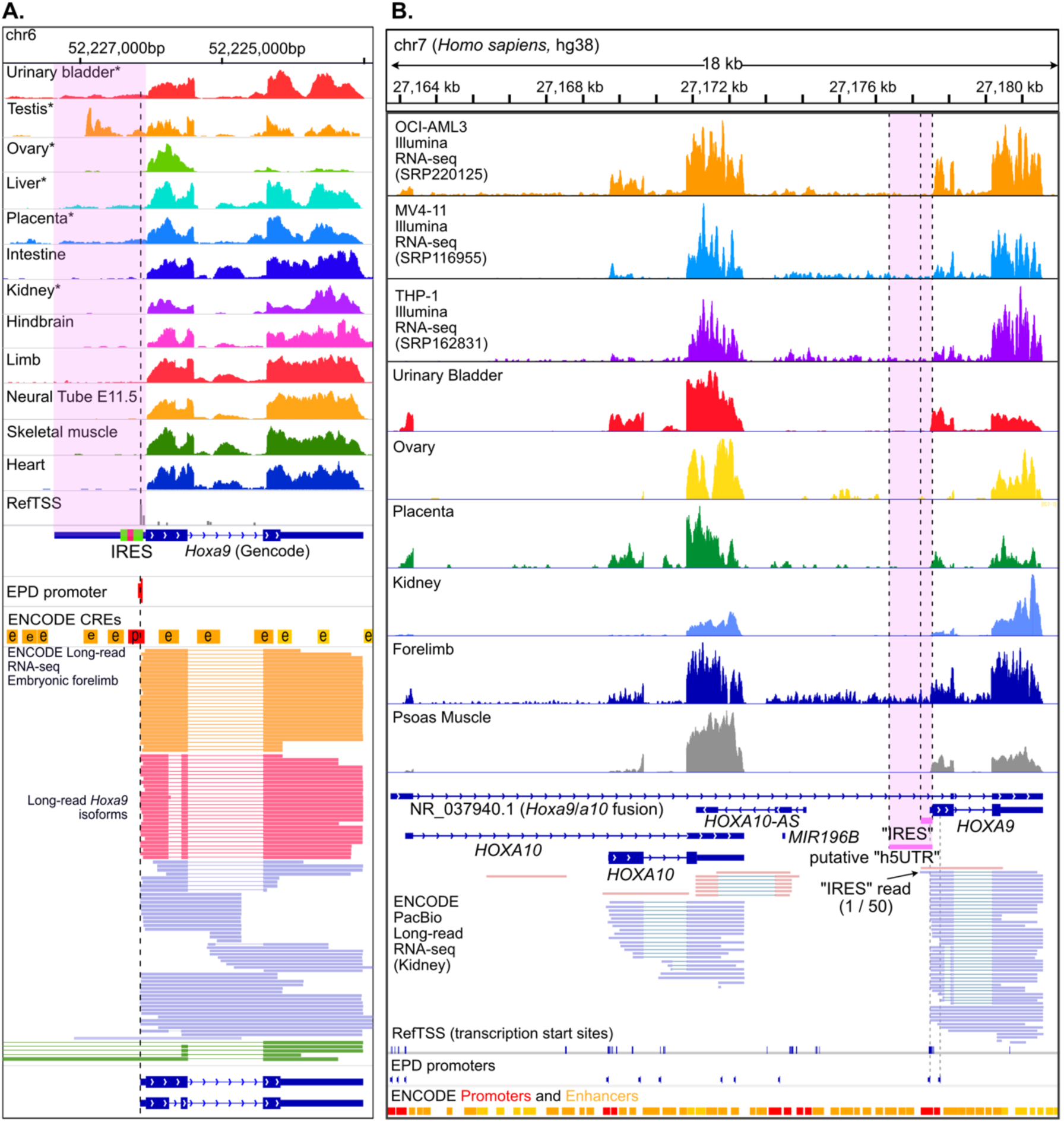
The putative IRES-like region of mouse and human *Hoxa9* are rarely expressed in mature transcripts. (A) ENCODE short-(upper) and long-read (PacBio, lower) RNA-seq data support transcription initiation almost exclusively downstream of the putative IRES-like region (shaded pink) in most mouse tissues, coinciding with an annotated promoter from EPD and ENCODE and 5’ CAGE data and refTSS sites. Long-read data additionally suggest the extended isoform annotation may reflect intronic RNA from *Hoxa9*/*a10* and *Mir196b*/*Hoxa9* fusion transcripts (green). Asterisks denote strand-specific RNA-seq. (B) Human *Hoxa9* expresses short 5’ UTR isoforms excluding the putative IRES. Genome browser tracks show short read polyA RNA-seq data from three Acute Myeloid Leukemia (AML) cell lines and data from the ENCODE project consortium from a variety of representative human tissues (upper). Long-read (PacBio Iso-seq) RNA-seq data from the ENCODE project (lower) shows two predominant isoforms whose transcripts initiate close to the *Hoxa9* protein coding sequence, consistent with annotated promoters (EPD and ENCODE) and transcription start sites (refTSS). One sense, and one antisense, Iso-seq read overlaps the putative IRES region.

**Figure S4.**
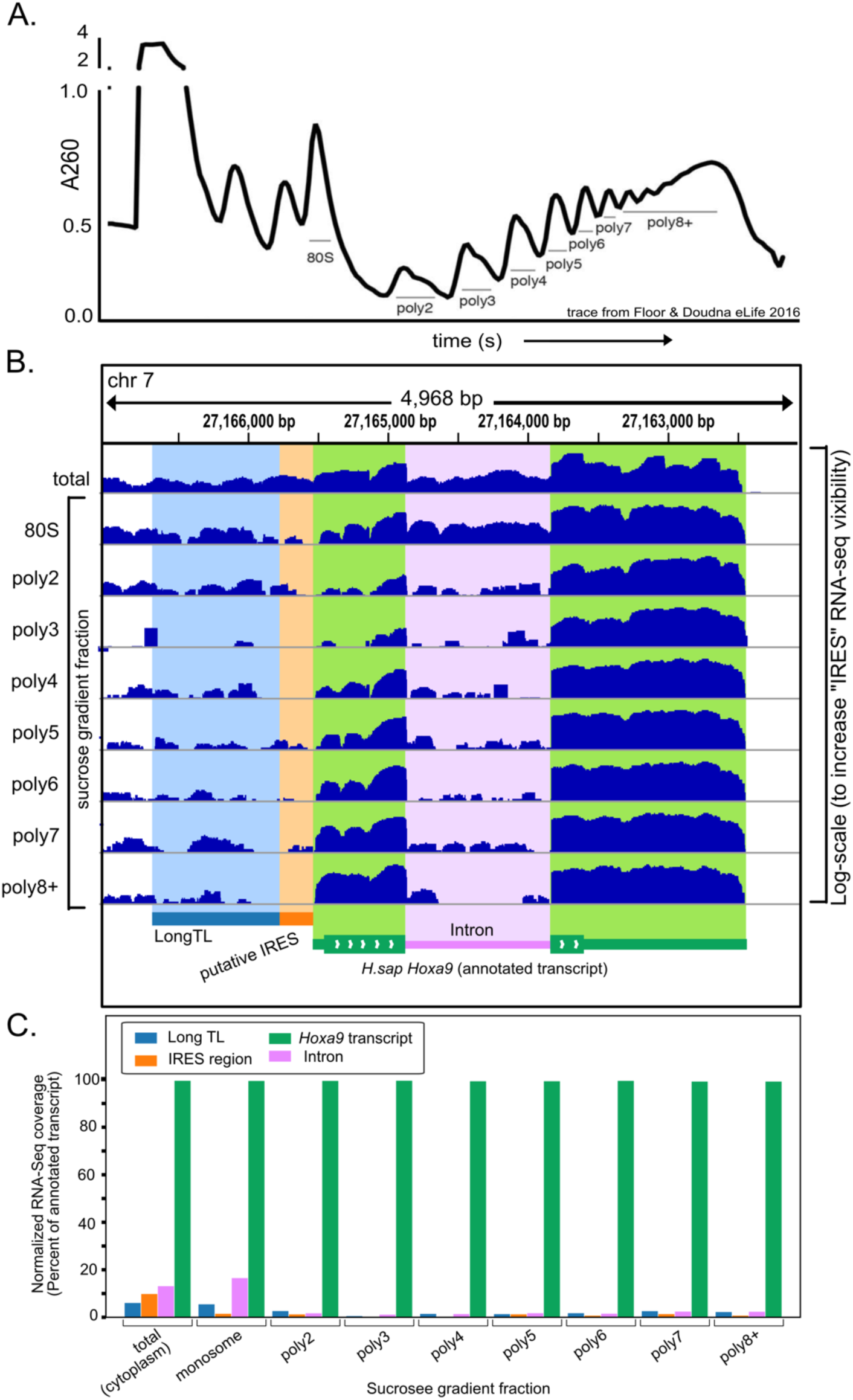
The extended 5’ UTR and putative IRES are absent from translating *Hoxa9* mRNA in HEK293T cells. A) Polysome extracts from HEK293T cells were separated by sucrose gradient fractionation and RNA-seq was performed (Floor and Doudna, 2015). (B) IGV browser image shows RNA-seq coverage over four regions around human *Hoxa9*. A log-scale was used to increase the visibility of coverage over the putative IRES region. (C) The average coverage over each region is plotted, normalized to coverage over the annotated transcript. The extended 5’ UTR and putative IRES regions have 5-10% the signal of the annotated transcript in total (ribo-depleted) RNA, which drops to ∼ 1% in polysomal fractions. Both putative IRES and intronic RNA are almost entirely absent from translating polysome fractions (poly2 and greater).

**Figure S5.**
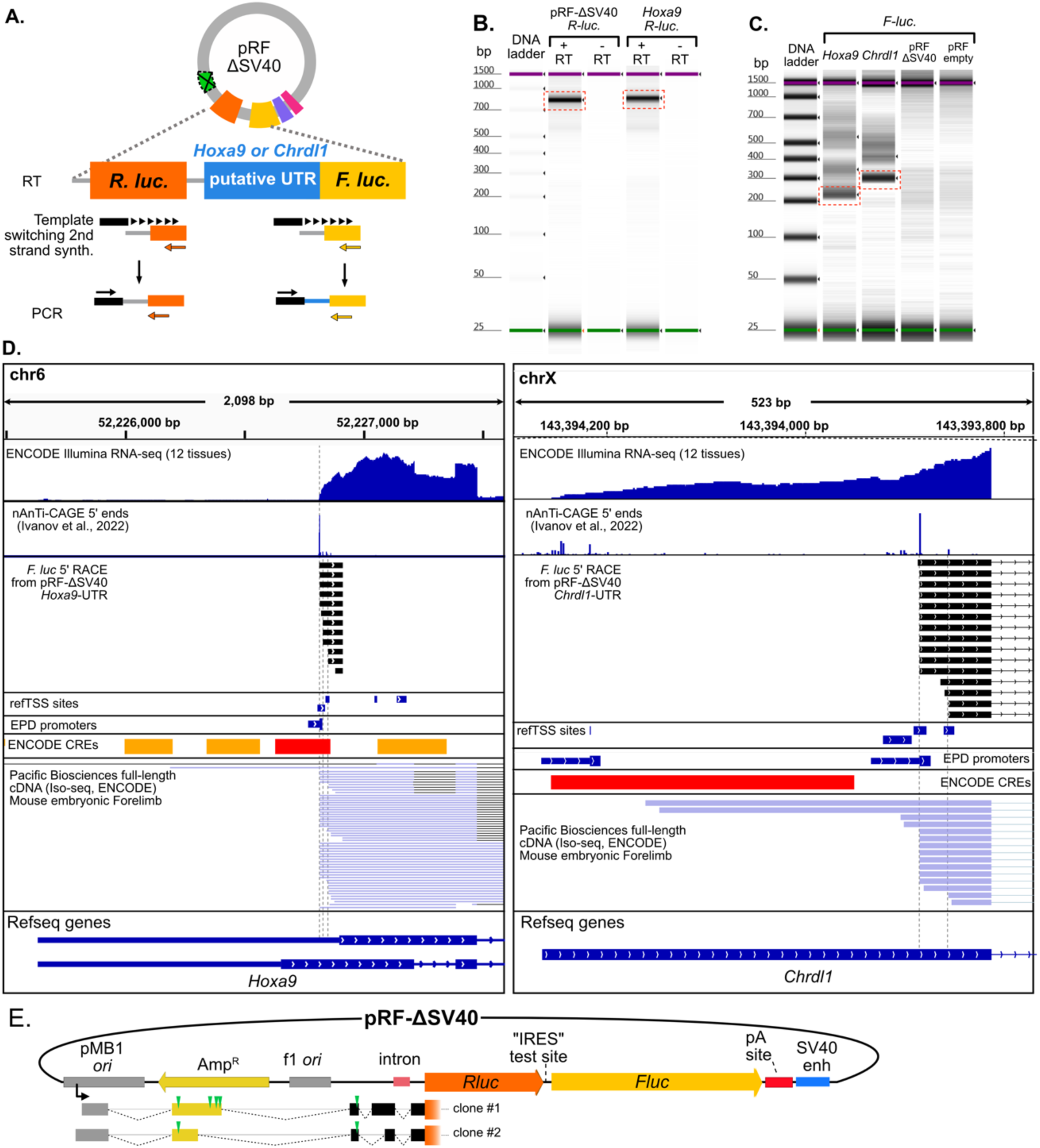
5’ RACE from pRF reporters recapitulates *in vivo* annotated *Hoxa9* and *Chrdl1*, TSS and identifies cryptic *Rluc* transcripts. (A) 5’ RACE was performed on *Rluc*. and *Fluc* transcripts from C3H10T1/2 cells transfected with *Hoxa9* and *Chrdl1* 5’ UTR pRF-ΔSV40 vectors. *Fluc* and *Rluc* RT and PCR primers are shown in yellow and orange, respectively, with the template switching oligo depicted in black. **(B)** *Rluc* RT-PCR products were electrophoresed on an Agilent TapeStation. A robust product (dotted red outlined box) was observed for both the *Hoxa9* 5’UTR and empty vector, despite the lack of an SV40 promotor. (**C)** *Fluc* RT-PCR electrophoresed as in B. Multiple short products (<700 bp) from *Hoxa9* and *Chrld1* putative UTRs, but not in empty-vector controls **(D)** The main RT-PCR products from *Hoxa9* and *Chrld1* transfections (C, red box) were cloned and sequenced. The cloned sequences, shown as genome browser tracks (black), recapitulate endogenous TSSs mapped with nAnTi-CAGE (Ivanov et al., 2022), and Iso-seq, from mouse embryonic tissues. **(E)** The RT-PCR products from *Rluc* (red box in B) were cloned and sequenced. Two clones are shown aligned to pRF-ΔSV40 with splice sites denoted with dotted black lines. These transcripts mapped to the pMB1 ori promoter (Lemp et al., 2012) and used many of the previously noted cryptic splice sites (Lemp et al., 2012). They are expected to have low translation efficiency due to multiple upstream AUG start codon uORFs (green triangles).

**Figure S6.**
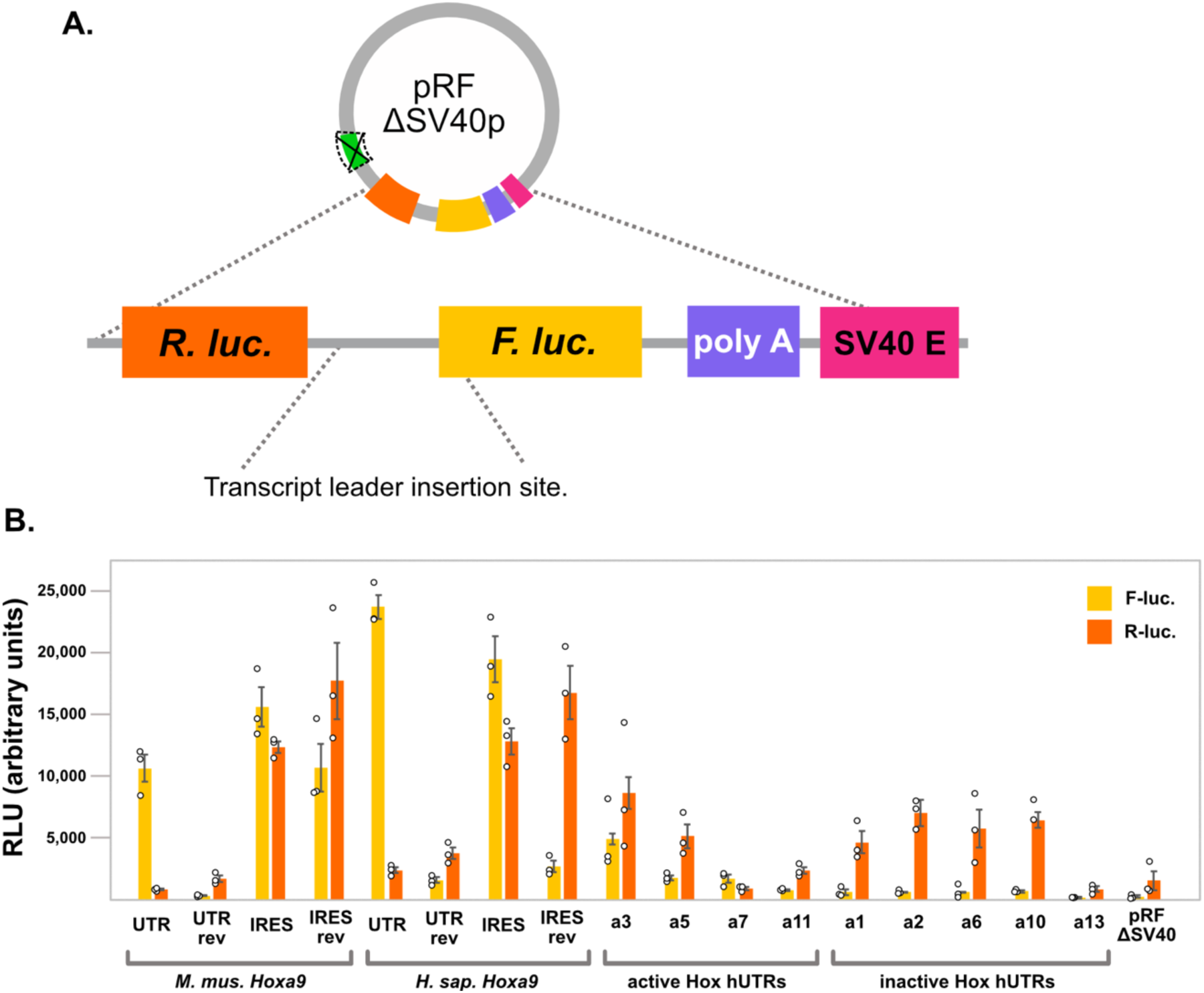
Annotated transcript leaders of *Hoxa* genes increase expression of both *Rluc* and *Fluc*. (A) Diagram of the “promoterless” bicistronic reporter plasmid pRF-ΔSV40. The putative IRES-like transcript leaders were cloned between *Renilla* (*Rluc*) and *Firefly* (*Fluc*) luciferase open reading frames and transfected into mouse mesenchymal cells. (B) Bar graphs showing raw luminescence values for *Rluc* and *Fluc* from each transfection. Error bars show standard error from three replicates. Most transcript leaders increase expression of both *Rluc* and *Fluc*, but to differing extents. For example, *M. mus. Hoxa9* UTR induces Fluc, but not *Rluc* expression well above background. The shorter “IRES-like” region induces expression of both, but Fluc is expressed more than *Rluc* compared to empty vector. The active (IRES-like) UTRs induce Fluc more than they induce *Rluc*, leading to a higher ratio (see figure 2), while the inactive UTRs induce similar fold changes in *Rluc* and *Fluc* expression, which the exception of *Hoxa13*, which induces neither gene. Note that this interpretation assumes does not account for potential variation in active plasmid concentrations during transfection. However, it seems unlikely that such variation could account for the variation in *Rluc* expression. Error bars show standard error with n = 3.

**Figure S7.**
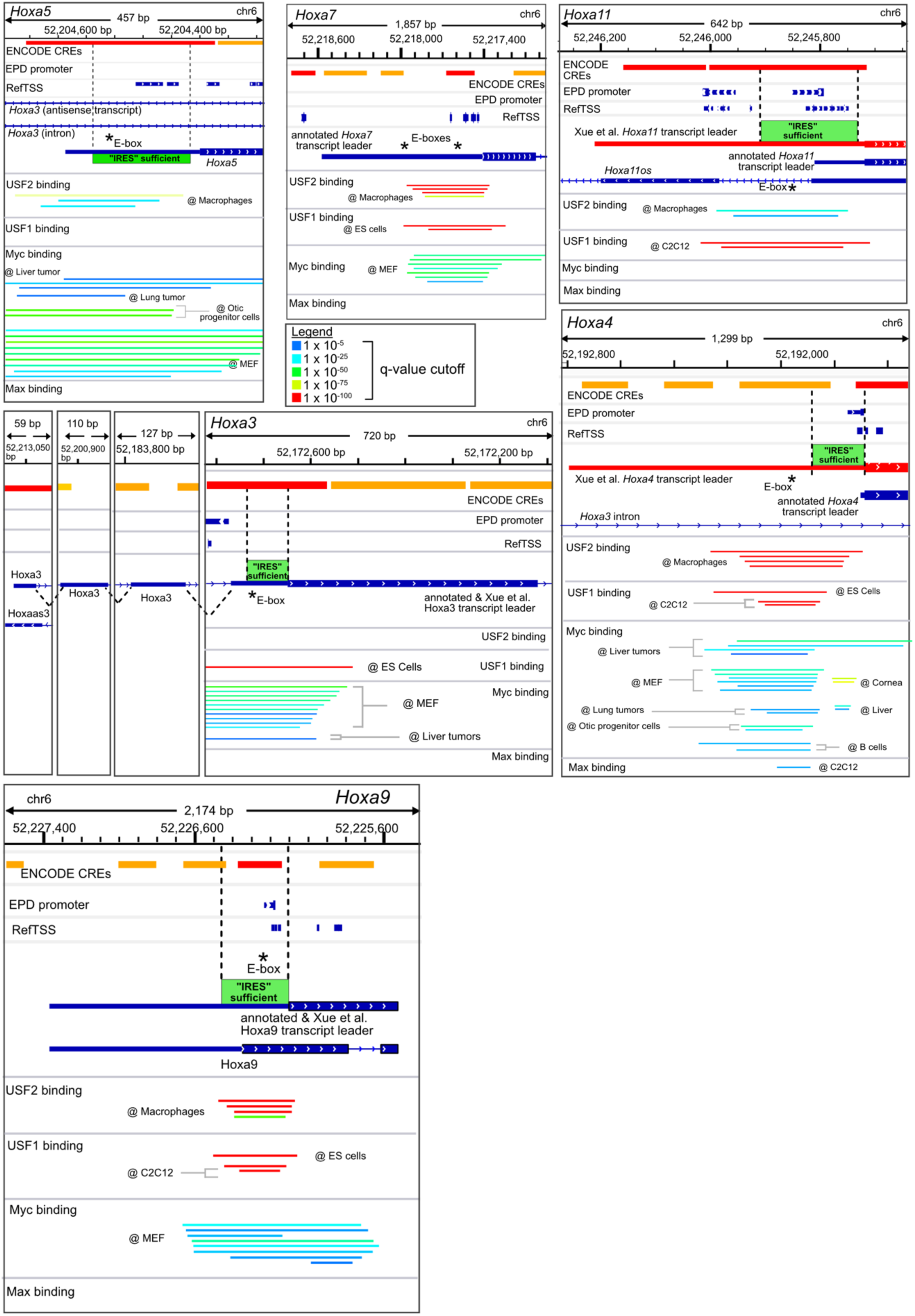
Mouse bHLH transcription factors bind to E-boxes near *Hoxa* gene promoters. Public mouse *USF1/2*, *MYC*, and *Max* ChIP-seq peaks were downloaded from chip-atlas.org (Oki et al., *EMBO* 2018). Colored lines depict sites of significant binding, (MACS2 (v 2.10), see legend for FDR). Regions previously found to be important for “IRES” like activity in bicistronic reporter assays are depicted as green boxes above the gene annotation models. These regions overlap promoter elements annotated by ENCODE and / or EPD, and most contain annotated transcription start sites from RefTSS. In all cases, each *Hoxa* gene promoter region shows binding by at least one of these factors, with particularly strong binding of *USF1* and *USF2* in the “P4” element from the putative IRES region of *Hoxa9*. This is consistent with ChIP-seq studies from human *Hoxa9* and provides further evidence supporting these sequences as E-boxes. *Hoxa* E-box binding was also seen for other bHLH TFs, including *TFE3* (*Hoxa3*, *a7,* and *a9*), *ARNTL* (*Hoxa3*, *a4*, *a7*, and *a9*), *BHLHE40*(*Hoxa7*), and *BHLHE41* (*Hoxa7*) (not shown).

**Figure S8.**
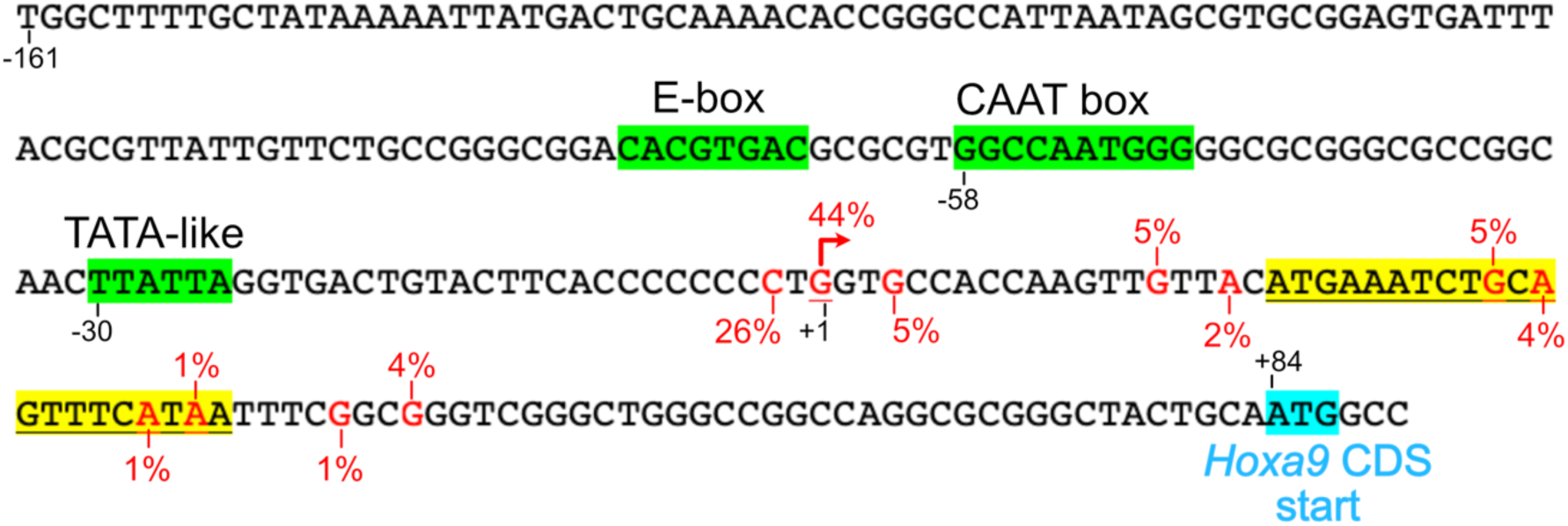
Sequence elements in the mouse *Hoxa9* promoter and 5’ transcript leader. Sequence above shows the region upstream of mouse *Hoxa9*. The E-box, CAAT box, and TATA like elements are highlighted in green. Transcription start sites are shown in red text, with the percentage of nAnTiCAGE reads mapping to each position in mouse E11.5 somites (Ivanov et al., 2022). The major TSS site is indicated with an arrow. A conserved uORF with a poor Kozak context (Ivanov et al., 2022) is highlighted in yellow. The *Hoxa9* CDS start codon is shown in blue.

**Figure S9.**
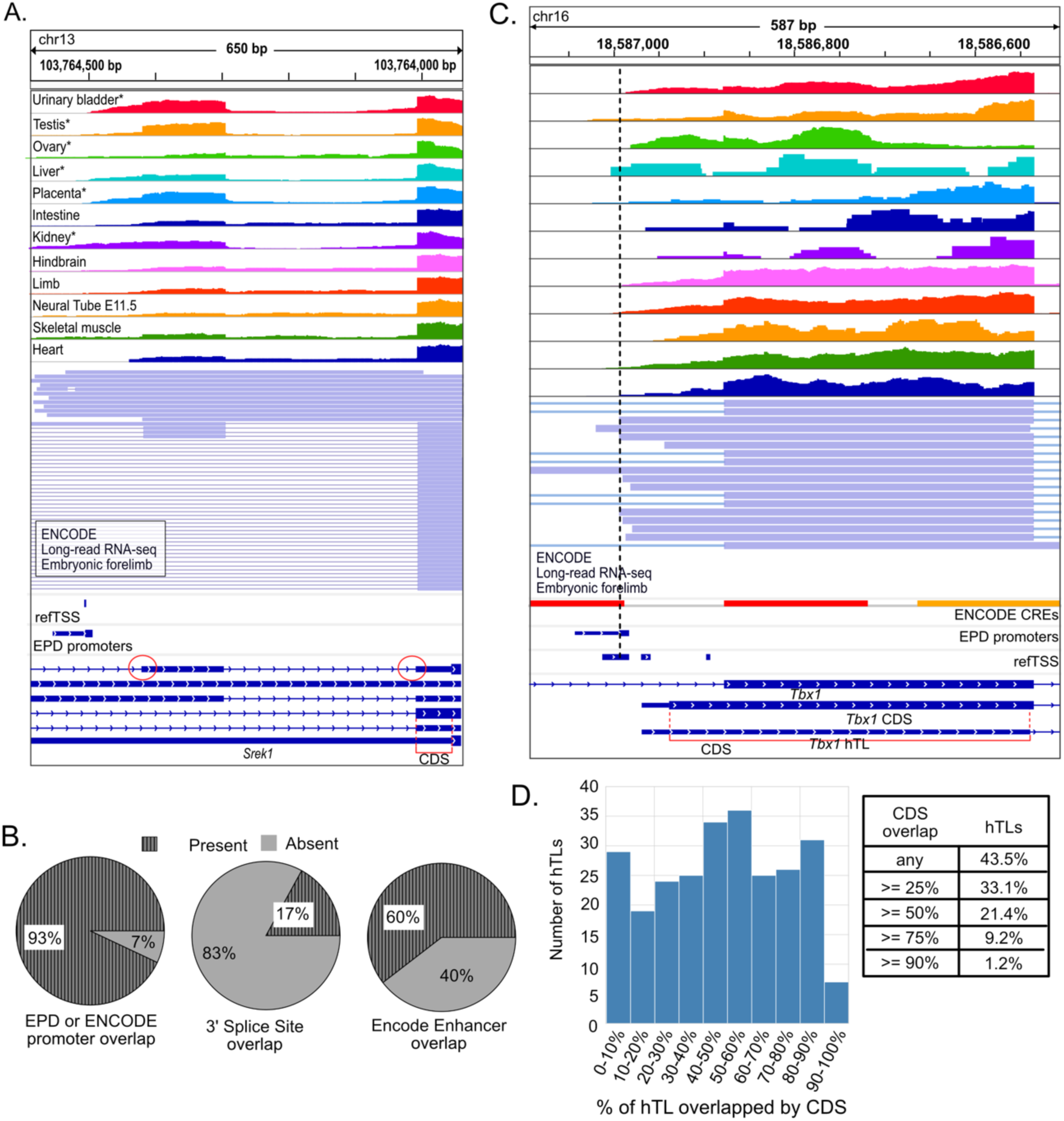
hTLs defined by Byeon et al. 2021 overlap promoters, splice sites, enhancers, and protein coding sequences. (A) Genome browser screenshot showing an example of a hTL from *Srek1*, which drives expression in the bicistronic reporter assay. Short- and Long-read RNA-seq suggest transcription initiates internally in this annotated transcript leader. The hTL overlaps an EPD promoter, two 3’ splice sites, and protein coding sequence from an alternatively spliced isoform (CDS) of the gene. (B) Pie graphs showing the percentage of hTLs that overlap EPD or ENCODE promoters, annotated 3’ splices sites, and ENCODE annotated transcriptional enhancer regions. (C) Genome browser screenshot showing a hTL from *Tbx1*, which almost entirely overlaps protein coding sequence from two other annotated transcript isoforms. The annotated protein coding sequence is translated in ribosome profiling data (GWIPs-VIZ; not shown) and has PhyloP conservation scores consistent with its translation (lower scores at wobble nucleotides). (D) Histogram showing the number of hTLs with varying percentages of CDS. 256 hTLs (43.5% of all hTLs) overlap annotated CDS regions, and a third of hTLs have at least 25% of their sequence overlapped with annotated CDS.

**Figure S10.**
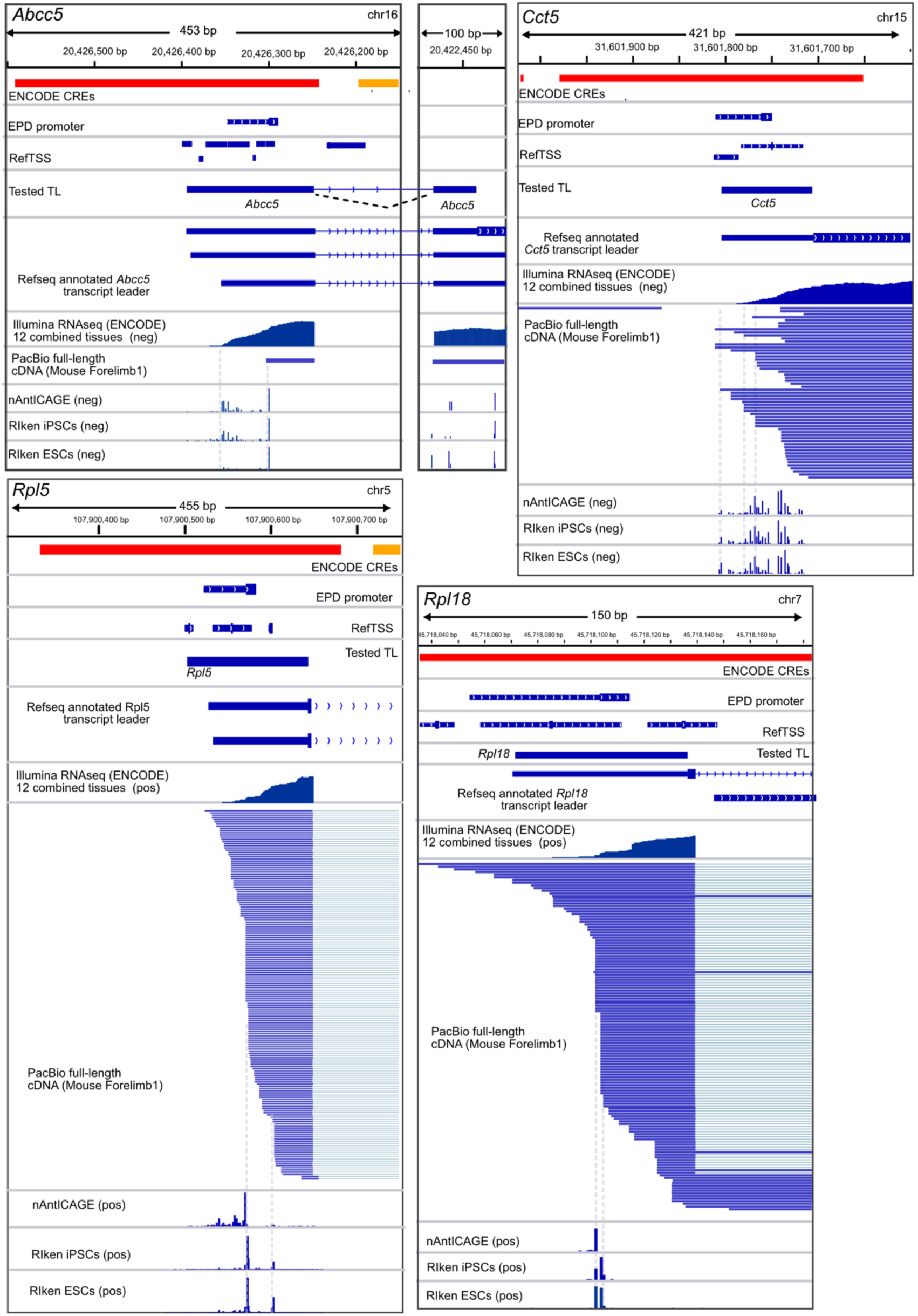
VELCRO-IP putative IRES elements contain annotated promoter sequences. Genome browser views show four putative IRESes from Leppek et al., 2021. EPD promoter elements are annotated within the putative IRES. PacBio Iso-seq and CAGE-seq data show numerous sites of internal initiation. Note, *Rpl5* was a “negative control”, as it did not interact with the ES9S helix *in vitro*, yet it had strong bicistronic reporter activity.

**Figure S11.**
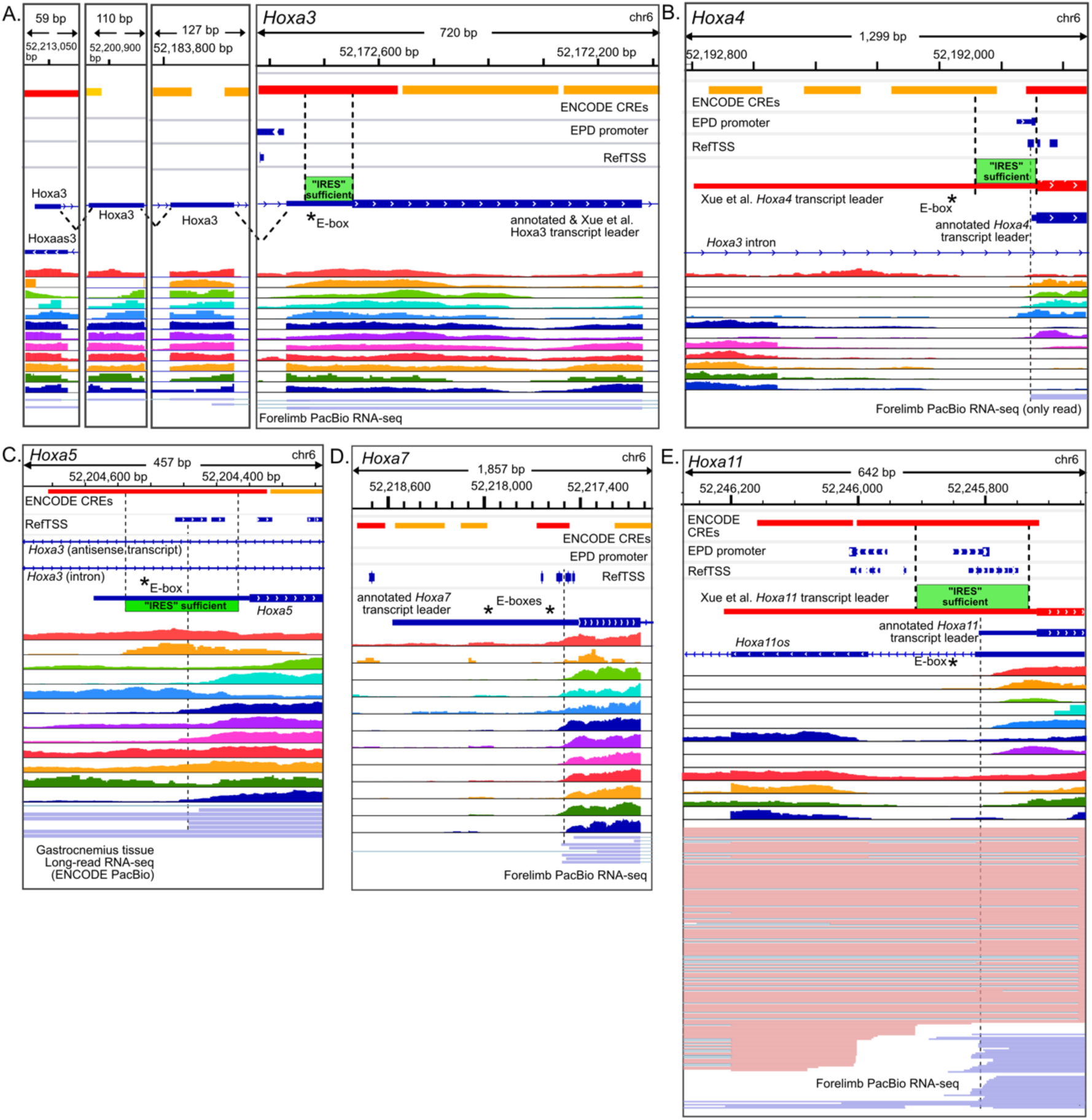
Misannotated transcript leaders and promoter overlap with putative IRESes from *Hoxa* genes. (A) The sufficient region of the putative *Hoxa3* IRES overlaps an ENCODE promoter. (B) The *Hoxa4* transcript leader mapped by Xue et al. 2015 is much longer than the annotated leader. The annotated, short leader is supported by short- and long-read RNA-seq data from ENCODE, and the sufficient region of the putative IRES overlaps an EPD promoter and refTSS sites. (C) The sufficient region of the putative *Hoxa5* IRES overlaps an ENCODE promoter and refTSS, which are supported by short- and long-read RNA-seq data. (D) The transcript leader of *Hoxa7* is much shorter than annotated, such that the region reported to be a putative IRES (Byeon et al., 2021) encompasses an ENCODE promoter and refTSS sites. (E) The *Hoxa11* transcript leader mapped by Xue et al. (2015) is much longer than the annotated transcript leader. The sufficient region of the putative IRES overlaps ENCODE and EPD promoters, and refTSS sites. Short- and long-read RNA-seq data support internal transcription initiation at the shorter annotated promoter. As with *Hoxa9* (Figure 2) extended, misannotated transcript leaders overlap introns in *Hoxa4*, and *Hoxa5*. * Asterisks show the locations of E-box motifs mutated in figure 3.

**Figure S12.**
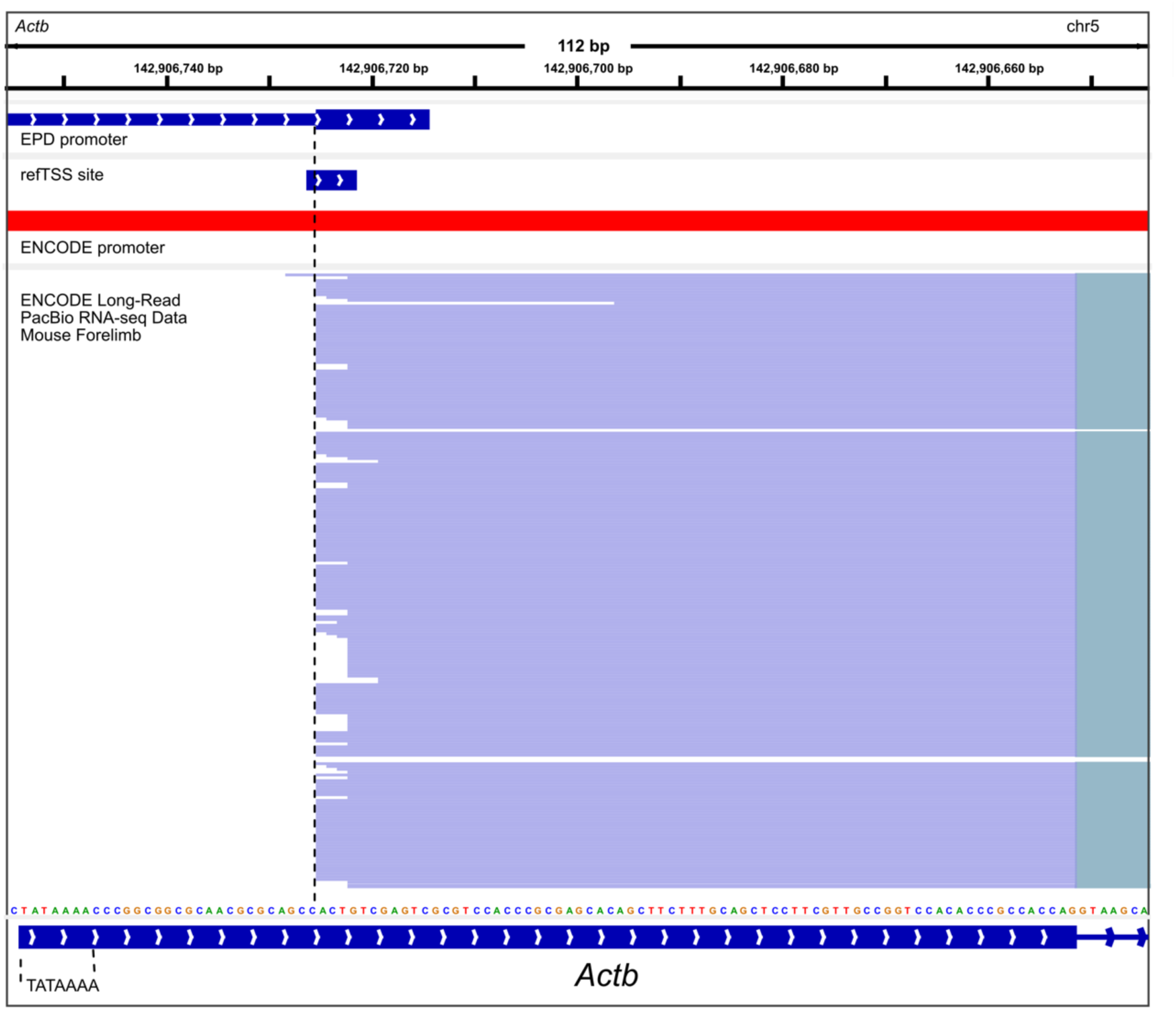
The transcript leader for mouse Beta actin is misannotated and includes a promoter. The refGene annotated transcript leader is shown. The annotated transcript leader begins with a TATA box, overlaps ENCODE and EPD promoters, and a refTSS site. Long-read RNA-seq data from ENCODE supports internal transcription initiation at the annotated promoter and refTSS site.

**Figure S13.**
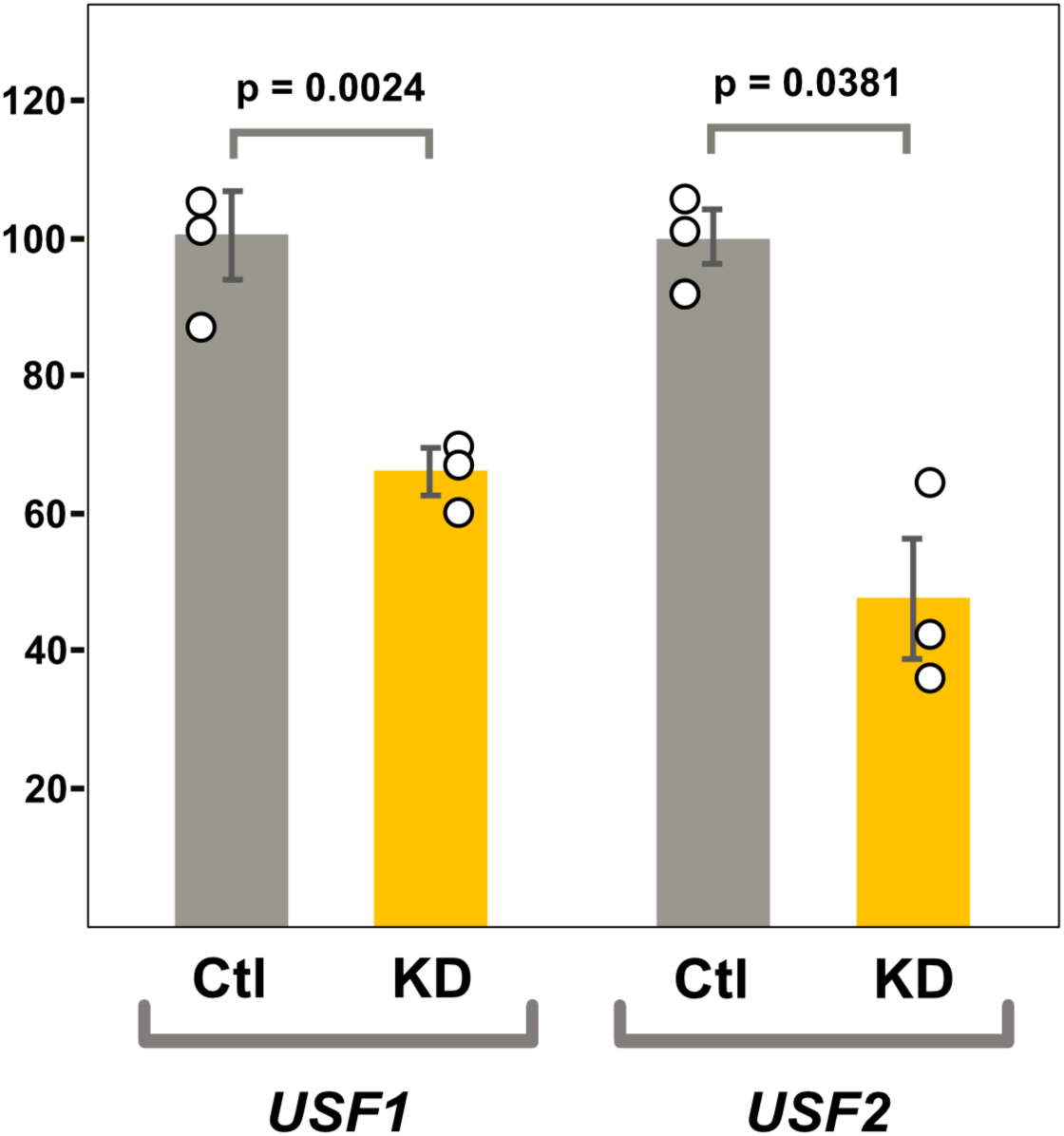
qPCR validation of *USF1*/*2* co-depletion by siRNA. *USF1* and *USF2* were assayed using qPCR to compare their mRNA levels from cells treated with control (scrambled) siRNA and cells treated with a mixture of *USF1* and *USF2* siRNA (Santa Cruz Biotechnology). Bar graphs show relative mRNA levels, normalized to control siRNA samples. Error bars indicate standard error. Both *USF1* and *USF2* mRNA levels were significantly depleted, compared to the scrambled control sample. P-values shown are from a 2-tailed, paired t-tests.

## Notes

### Competing Interest Statement

The authors have declared no competing interest.

### Summary of Updates

In this revision, we added new 5' RACE analyses of transcripts originating from pRF-DeltaSV40 plasmids (new Figure S5), which support the presence of promoters in the putative Hoxa9 and Chrdl1 IRESes and provide a rationale for the varying Rluc expression seen from our reporters. We also added new public ChIP-seq data that show binding of USF1, USF2, MYC, and MAX at the mouse Hoxa gene E-boxes (new Figure S7), and RNA polymerase II ChIP-seq data consistent with the putative Hoxa9 IRES being a promoter (new Figure 2). In response to public comments on our first preprint, we also updated Figure 2 (which is now split into Figures 2 and 3), expanded supplemental Figure S3, added new supplemental Figures S4 and S10, and revised some language for clarity. We also expanded the discussion section to address these added results.

